# A family portrait of the genomic factors shaping tandem repeat mutagenesis

**DOI:** 10.64898/2026.03.06.710071

**Authors:** Thomas A. Sasani, Michael E. Goldberg, Akshay K. Avvaru, Thomas J. Nicholas, Deborah W. Neklason, Egor Dolzhenko, Tom Mokveld, Katherine M. Munson, Kendra Hoekzema, Marcelo Ayllon, Eli J. Kaufman, David Porubsky, Zev Kronenberg, Guilherme de Sena Brandine, William J. Rowell, Lynn B. Jorde, Christopher E. Mason, Michael A. Eberle, Paul N. Valdmanis, Evan E. Eichler, Aaron R. Quinlan, Harriet Dashnow

**Affiliations:** Department of Human Genetics, University of Utah, Salt Lake City, UT, USA; Department of Biomedical Informatics, University of Colorado Anschutz, Aurora, CO, USA; Department of Internal Medicine, University of Utah, Salt Lake City, UT, USA; PacBio, Menlo Park, California, USA; Department of Genome Sciences, University of Washington School of Medicine, Seattle, WA, USA; Division of Medical Genetics, University of Washington School of Medicine, Seattle, WA, USA; European Molecular Biology Laboratory (EMBL), Genome Biology Unit, Heidelberg, Germany; Department of Systems and Computational Biomedicine, Weill Cornell Medicine, New York, NY, USA; Trivedi Institute of Space and Global Biomedicine, University of Pittsburgh, Pittsburgh, PA, USA; Howard Hughes Medical Institute, University of Washington, Seattle, WA, USA

## Abstract

Tandem repeats (TRs) are among the most mutable loci in the human genome, but the genomic determinants of TR mutagenesis remain mysterious. We used PacBio HiFi long-read sequencing to profile nearly eight million TR loci in 28 members of a large, four-generation CEPH/Utah family designated K1463. We identified 1,270 *de novo* TR expansions and contractions across 20 children in the pedigree. *De novo* mutations (DNMs) were more likely to occur at loci that were longer, composed of uninterrupted motif sequences, and heterozygous in the parental germline. Children born to older fathers also exhibited more *de novo* mutations at short tandem repeats (STRs). A total of 43 TR loci were hyper-mutable in K1463, expanding or contracting up to twelve times across the pedigree. Among hyper-mutable loci that comprised multiple motifs (i.e., “complex” loci), specific motifs expanded and contracted more often than others; for example, all ten DNMs at a complex, hyper-mutable locus near the non-coding RNA *LINC03021* involved the same 19bp motif. The mutability of particular motifs may be attributable to allele length, as 95% of DNMs at complex loci were expansions and contractions of the most abundant motif on a parental haplotype. However, future work will be required to disentangle the effects of nucleotide content and allele length on motif-specific mutability, especially at hyper-mutable TRs. Overall, this study combines long-read sequencing technologies with new software tools to comprehensively investigate the factors that influence TR mutagenesis.

## Introduction

The human genome is replete with repetitive arrays of nucleotide motifs called tandem repeats (TRs). TRs are typically classified as either short tandem repeats (STRs), composed of 1-6 base pair motifs, or variable number of tandem repeats (VNTRs), which contain 7+ base pair motifs. Expansions and contractions at more than 70 TR loci underlie monogenic disorders, including Huntington’s disease, certain forms of spinocerebellar ataxia, and adult familial myoclonic epilepsy [1–4]. TR polymorphisms are also associated with complex, polygenic traits, including human height [5,6], lipoprotein A concentration, male pattern baldness, and serum urea concentration [6].

*De novo* mutations (DNMs) at TRs are often caused by a phenomenon called “slipped-strand mispairing,” in which newly synthesized nucleotides on the nascent strand are improperly aligned to the template during DNA replication [7,8]. Slipped-strand mispairing is likely responsible for the majority of *de novo* mutations at STRs [8–10], though unequal crossing over between homologous chromosomes may generate a smaller number of *de novo* mutations at both STRs and VNTRs [7,9–15]. We previously observed that mutation rates decrease as a function of motif size at STRs but increase as a function of VNTR motif size [16], reflecting a potential transition point between strand slippage and unequal crossing over as the dominant mechanisms underlying TR expansions and contractions.

DNMs at TR loci are typically detected by genotyping family members at a predefined list of loci in the human genome. Most studies have estimated the germline mutation rate at STRs to be between 4.95 x 10^-5^ and 5.6 x 10^-5^ per locus per generation, over three orders of magnitude higher than at single-nucleotide sites [17–22]. As short-read sequencing technologies cannot reliably genotype large VNTR alleles, direct estimates of the human germline VNTR mutation rate are limited, though VNTRs are known to be highly polymorphic in human populations [23,24]. Using long PacBio HiFi sequencing reads, we recently estimated the STR and VNTR mutation rates to be 1.1 x 10^-5^ and 1.7 x 10^-6^ per locus, per generation, respectively [16]. Since TR mutation rates are calculated by dividing an observed number of *de novo* alleles by the total number of genotyped reference loci, the methods used to construct those reference loci catalogs can have a dramatic impact on mutation rate estimates. We likely estimated a lower STR mutation rate because we searched for *de novo* mutations across a much larger catalog of TR loci than previous studies, and because we did not require those loci to be polymorphic in the cohort.

Germline mutation rates at STR loci covary with numerous genomic factors. Dinucleotide motifs are the most mutable STRs, and STR mutation rates generally decline as a function of motif size [16–18,25]. “Pure” STR alleles, comprising uninterrupted arrays of a particular *k-*mer nucleotide motif, are more likely to mutate than “interrupted” alleles [26–30]. Longer STR alleles also tend to be more polymorphic and exhibit higher mutation rates than shorter alleles [16,31]. The sequence composition of individual motifs likely influences STR mutation rates, as well; `AT` and `AA[A/G]G` repeats are some of the most mutable dinucleotide and tetranucleotide motifs, respectively [10,20,32,33]. On *Escherichia coli* plasmid sequences and in human cell lines, transcribed STR alleles exhibit greater instability than their untranscribed complements, possibly due to collisions between the replication fork and transcriptional machinery [34–36]. Consistent with a replicative origin for *de novo* TR mutations, fathers contribute more *de novo* STR alleles than mothers, likely owing to the continuous mitotic divisions that occur in spermatogonial stem cells [16,17,19,26,37,38].

The genomic determinants of mutation rates at VNTR loci are more mysterious. In both humans and *Saccharomyces cerevisiae*, VNTR allele size is strongly correlated with mutation rate [24,39,40]. Certain VNTR loci, at which >1% of sequenced offspring harbor a *de novo* allele, have been reported to be “hyper-mutable” in human genomes [24]. A hyper-mutable VNTR locus was previously identified in multiple CEPH/Utah pedigrees [41], at which 52 of 310 children harbored a *de novo* expansion or contraction. At another hyper-mutable VNTR locus, larger alleles were more prone to *de novo* mutations and produced smaller *de novo* allele changes more often than large ones [40]. VNTRs comprise larger motifs than STRs, and VNTR loci are often larger than an average Illumina sequencing read. As a result, VNTR loci are more difficult to profile using short-read sequencing technologies than STRs.

We recently used PacBio HiFi and Element AVITI sequencing data to identify *de novo* tandem repeat mutations in a large CEPH/Utah family designated K1463 [16,42]. Here, we substantially expand our analysis of TR mutagenesis in K1463 by augmenting the sequencing depth of key family members and extending our analysis to a fourth generation. By resequencing the parents of twelve K1463 children, we nearly doubled our callset of *de novo* TR mutations. We evaluated the mutability of specific STR motifs, as well as the effects of allele length, purity, and parental heterozygosity on TR mutagenesis. We also found that the number of STR DNMs increases with paternal age. Finally, we carefully analyzed the motif composition of many hyper-mutable tandem repeat loci, including a single locus that mutated 10 times in K1463. At this locus, we found that two VNTR motifs — which differ by only two base pairs — exhibited substantially different mutation rates in the K1463 pedigree. However, we also observed that allele length was highly correlated with motif-specific mutability at complex TR loci, raising the possibility that subtle differences in motif composition, allele length, or a combination of the two, have substantial effects on TR mutagenesis.

## Results

### Identifying de novo expansions and contractions at TR loci

We previously created a comprehensive catalog of approximately 7.8 million tandem repeat (TR) loci in the human genome [16]. This catalog included every TR locus in the T2T/CHM13 reference genome that measured between 10 and 10,000 base pairs; if two or more TR loci were within 50 base pairs of each other, we merged them into a single locus (**Figure 1a**). We additionally classified all TR loci as either “simple” or “complex” according to their motif composition. “Simple” TR loci comprised a single motif, while “complex” TR loci comprised two or more unique motifs (e.g. [CAG]_n_[ATTGGGG]_n_) (**Fig. 1a)**. We used TRGT [43] to genotype each member of the K1463 pedigree at these 7.8M loci. We then used TRGT-denovo [44] to identify *de novo* tandem repeat alleles in each trio.

**Figure 1:**
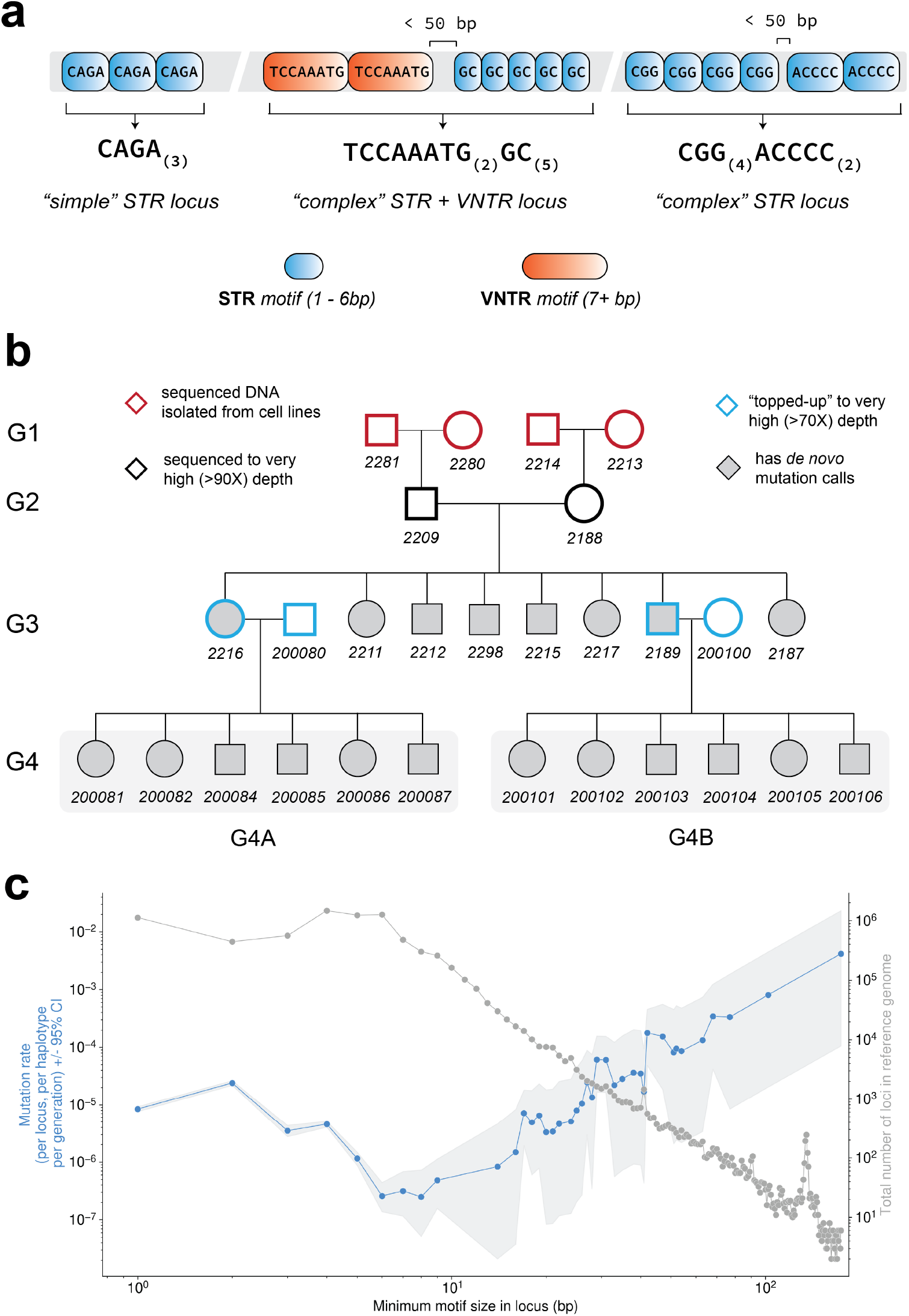
Identifying *de novo* tandem repeat expansions and contractions in the CEPH K1463 pedigree. **a)** We previously created a catalog of approximately 7.8 million tandem repeat (TR) loci [16] using Tandem Repeats Finder [47] and TRSolve [43]. We identified all TR sequences measuring between 10 and 10,000 base pairs in the T2T/CHM13 reference assembly; if two TR sequences were within 50 bp of each other in the reference genome, they were merged into a single TR locus. If the nearby loci comprised more than one unique motif, we described the merged locus as “complex.” **b)** K1463 comprises four generations (G1 through G4). As described in a previous manuscript [16], we sequenced blood-derived DNA from the members of G2-4 (and cell line DNA from the members of G1) using the PacBio HiFi platform. The members of G2 were sequenced to an average of 90-100X depth, the parents of G4 were “topped up” to approximately 70X (see **Supplementary Note**), and all other family members were sequenced to an average of 40X. **c)** We used TRGT [43] to genotype every member of the CEPH/Utah K1463 pedigree at each TR locus, and used TRGT-denovo [44] to identify *de novo* expansions and contractions of TR alleles in each trio from G3 and G4. We calculated the mutation rate (expressed per locus, per haplotype, per generation) for all loci (shown in blue), stratified by the size of the smallest motif within the locus. The 95% confidence interval (calculated using a Chi-square approximation) is shown for each mutation rate estimate as a light grey band. The average number of loci per child that passed filtering criteria are shown as connected grey points.

### Extending our analysis of de novo TR alleles to the fourth generation of K1463

Our prior study of *de novo* TR mutations in K1463 was limited to DNMs observed in the third generation (G3) for two primary reasons: first, the sequencing data from G1 individuals were derived from lymphoblastoid cell lines rather than whole blood; and second, we observed a high number of false positive *de novo* alleles in G4, which we attributed to allelic dropout in a parent (see **Supplementary Note** for additional details) [16]. To address challenges associated with allelic dropout in this study, we performed “top-up” sequencing on two members of G3 and each of their spouses (**Materials and Methods**; **Fig. 1b**). We then re-genotyped every member of the K1463 pedigree and identified *de novo* mutations using updated versions of TRGT and TRGT-denovo, producing a comprehensive new K1463 DNM callset. We also re-performed small variant calling and haplotype phasing with DeepVariant and HiPhase [45,46] (**Materials and Methods**). In our updated callset, we identified 1,270 TR *de novo* mutations in G3 and G4. Of these, 805 DNMs (63%) occurred at complex loci comprising more than one unique tandem repeat motif.

### Long reads enable precise de novo size estimation

We can typically estimate the size of a *de novo* TR expansion or contraction by comparing the *de novo* allele length to the lengths of the diploid TR alleles in both parents. If the *de novo* TR allele can be assigned to a parent-of-origin (PO), we can improve our estimate by comparing the *de novo* allele length to the TR allele lengths in the PO. In either approach, the smallest allele length difference is considered the most parsimonious mutational change. Using short read sequencing data from two-generation trios, however, it is generally not possible to infer the specific parental *allele* on which the *de novo* TR mutation occurred (which we refer to as the “precursor” allele). By applying HiPhase [46] to long-read PacBio HiFi sequencing, we could infer both the parent-of-origin and the exact “precursor” parental allele that mutated at 688 TR DNMs, enabling size estimation of expansions and contractions at these loci. The parsimony-based estimate disagreed with the precursor-based estimate at 23% (161 / 688) of DNMs for which we could apply both approaches (**Supplementary Figure 1**), demonstrating the benefit of phased, long-read sequencing data for measuring expansion and contractions. Compared to DNMs with concordant size estimates, the DNMs with conflicting size estimates contained similar proportions of expansions and contractions (Chi-square contingency *p* = 0.64). At one example TR locus, the parsimony-based approach suggested that a *de novo* allele was a 50 base pair contraction, while an exact comparison to the precursor allele demonstrated that it was, in fact, a 75 base pair *expansion*. For all subsequent analyses, we considered the phased precursor-based size estimate to be correct if the two approaches disagreed, though we used parsimony-based size estimates if the precursor allele could not be inferred.

### Most TR DNMs are small expansions and contractions

We observed similar numbers of expansions and contractions across our TR DNM callset (629 expansions and 641 contractions, binomial *p =* 0.379). Approximately one-third of DNMs (*n =* 465) DNMs occurred at simple TR loci, making it trivial to determine the motif that expanded or contracted to produce a *de novo* allele. For the remaining 805 TR DNMs, which occurred at complex loci comprising multiple unique motifs, it was sometimes difficult to determine the exact motif that expanded or contracted. However, by inferring the parental “precursor” allele at a subset of these complex DNMs, we could directly compare the child’s *de novo* allele sequence to the precursor allele sequence and identify the specific motif that mutated (see **Materials and Methods)**. In total, there were 586 DNMs that either a) occurred at simple TR loci, or b) occurred at complex TR loci for which we could accurately determine the specific motif that expanded or contracted. Among these 586 DNMs, non-homopolymer STR DNMs were enriched for contractions (164 *de novo* alleles were expansions and 202 were contractions; binomial *p* = 2.6 x 10^-2^), though this enrichment was almost exclusively attributable to dinucleotide motifs (61 expansions and 102 contractions, binomial *p* = 8 x 10^-4^, **Fig. 2**). Nearly all prior studies of genome-wide germline *de novo* STR mutations reported a bias toward expansions [17–19]. Each of these studies used short-read sequencing data to genotype smaller catalogs of STRs, raising the possibility that differences in sequencing technologies, genotyping methods, and/or the composition of TR loci catalogs could be responsible for our observed lack of an expansion bias. To address the first of these hypotheses, we genotyped a smaller collection of tandem repeat loci — which was previously profiled using short-read sequencing data by Mousavi et al. [48] — using our long-read sequencing pipeline. This previously published catalog comprises STR loci that are shorter, on average, than loci in our catalog; the TR locus N50 in our catalog was 63 bp, while the N50 in [48] was 15 bp (**Supp. Fig. 2**). After genotyping this orthogonal catalog, we found a similar enrichment of contractions over expansions at dinucleotide motifs (binomial *p* = 1.4 x 10^-2^; **Supp. Fig. 3**; **Materials and Methods**), and no bias toward expansions for any motif size. Despite their differences in TR composition, we did not observe an enrichment of expansions using either catalog, suggesting that technical differences between short- and long-read sequencing technologies and/or genotyping approaches are likely responsible for the discordance between our results and previous *de novo* TR size estimates. Notably, almost half of the *de novo* alleles we identified in K1463 (*n* = 620) were larger than 150 base pairs and would be very challenging to identify using short reads alone.

**Figure 2:**
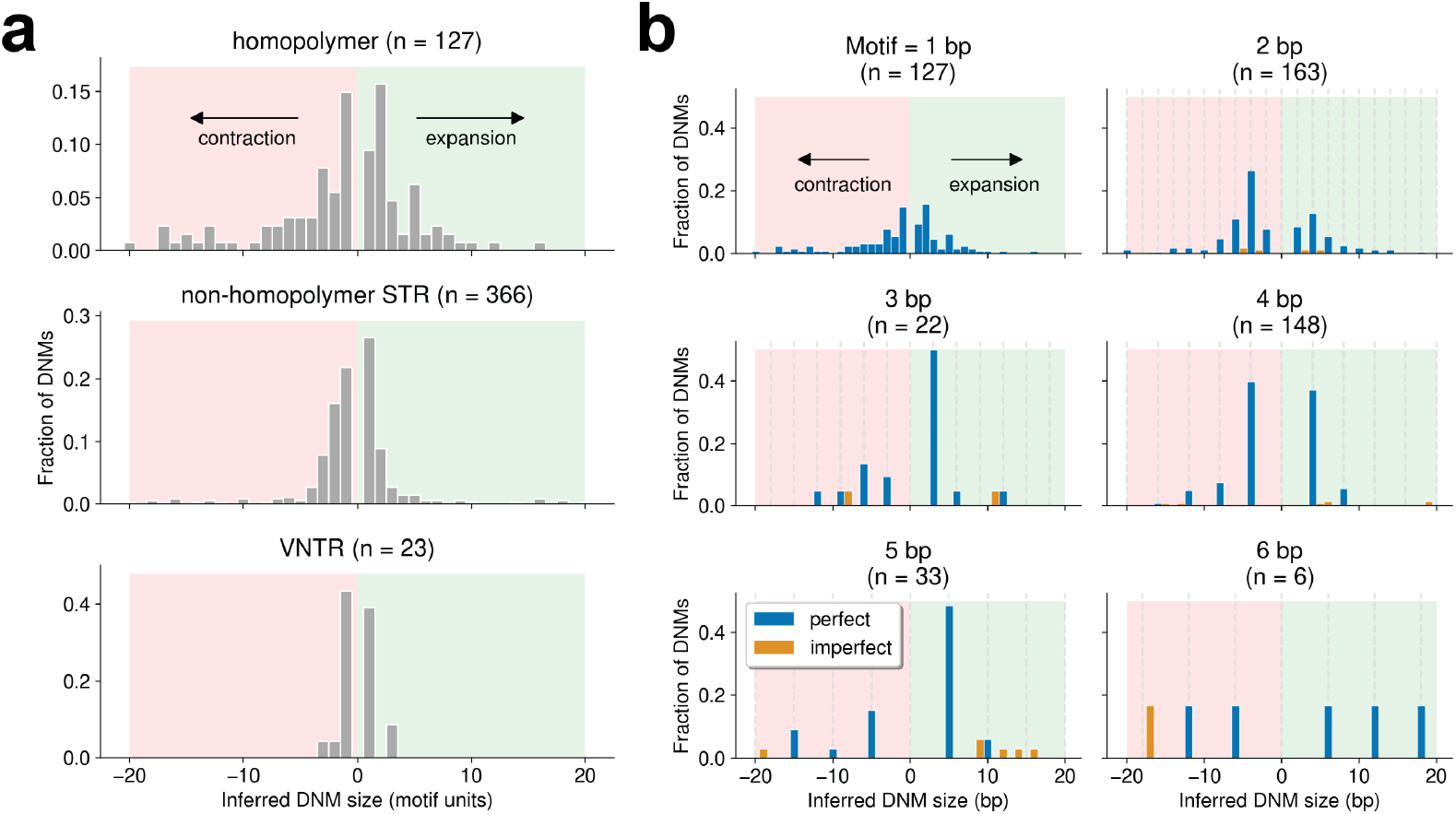
Most *de novo* mutations are small expansions and contractions at STRs. **a)** Number of TR DNMs that involved an expansion or contraction of the specified number of repeat units: homopolymer (1 bp), non-homopolymer STR (2-6 bp) and VNTR (7+ bp). These plots only include DNMs at which we could determine the motif that mutated (*n* = 586 loci that comprised a single motif or comprised multiple motifs that could be fully phased and decomposed using a post-hoc method; see **Materials and Methods**). **b)** Number of STR DNMs that involved an expansion or contraction of the specified number of base pairs, stratified by the size of the motif that mutated at the locus. Blue bars correspond to counts of DNMs at which the *de novo* allele size was perfectly divisible by the motif size (e.g., an 8bp *de novo* allele at a locus composed of 4bp motif), and orange bars correspond to DNMs that were not perfectly divisible (e.g., a 9bp expansion of a 4bp motif). Dotted vertical lines indicate the expected sizes of perfectly divisible expansions and contractions.

### De novo expansions and contractions are more likely to occur at heterozygous loci, and on long and pure alleles

Although we could only infer the parental “precursor” allele for 688 TR DNMs in the K1463 pedigree, we used HiPhase [46] haplotype assignments to infer the parent-of-origin for 904 *de novo* TR alleles (**Materials and Methods**). At every *de novo* mutation with an inferred parent-of-origin, we compared TR genotypes between the parent-of-origin (PO) and the other parent (*not* the parent-of-origin; NPO). At loci composed of STR motifs, we observed that the parent in whom the TR DNM occurred was more likely to be heterozygous than the other parent (Chi-square *p* = 4.1 x 10^-3^; **Fig. 3a-b**). By identifying the specific parental “precursor” allele that mutated to produce a *de novo* expansion or contraction, we also compared various features of the TR alleles that did or did not mutate in a parent’s germline. For example, within the PO’s germline, precursor TR alleles were longer than the alleles on homologous chromosomes (Wilcoxon signed-rank *p* = 4.53 x 10^-15^; **Fig. 3c**). Similarly, TR alleles that expanded or contracted were purer than their homologous alleles (Wilcoxon signed-rank *p* = 4.4 x 10^-13^; **Fig. 3d**). Although prior studies have demonstrated that longer and purer TR alleles are more prone to mutation, we were able to identify the specific parental allele that underwent a *de novo* expansion or contraction, enabling us to precisely measure the characteristics of these precursor alleles. We also searched for evidence that TR loci with *de novo* mutations in K1463 were more mutable in an orthogonal human genome dataset. At every locus with a *de novo* mutation in K1463 (*n* = 1,270 loci), we genotyped 100 members of the Human Pangenome References Consortium (HPRC) [49] and calculated the fraction of HPRC individuals who possessed two non-identical TR alleles; we refer to this measure as “locus heterozygosity.” Compared to TR loci that did not mutate in K1463, HPRC locus heterozygosity was consistently higher at loci with *de novo* expansions or contractions, regardless of motif composition at those loci (**Supp. Fig. 4**).

**Figure 3:**
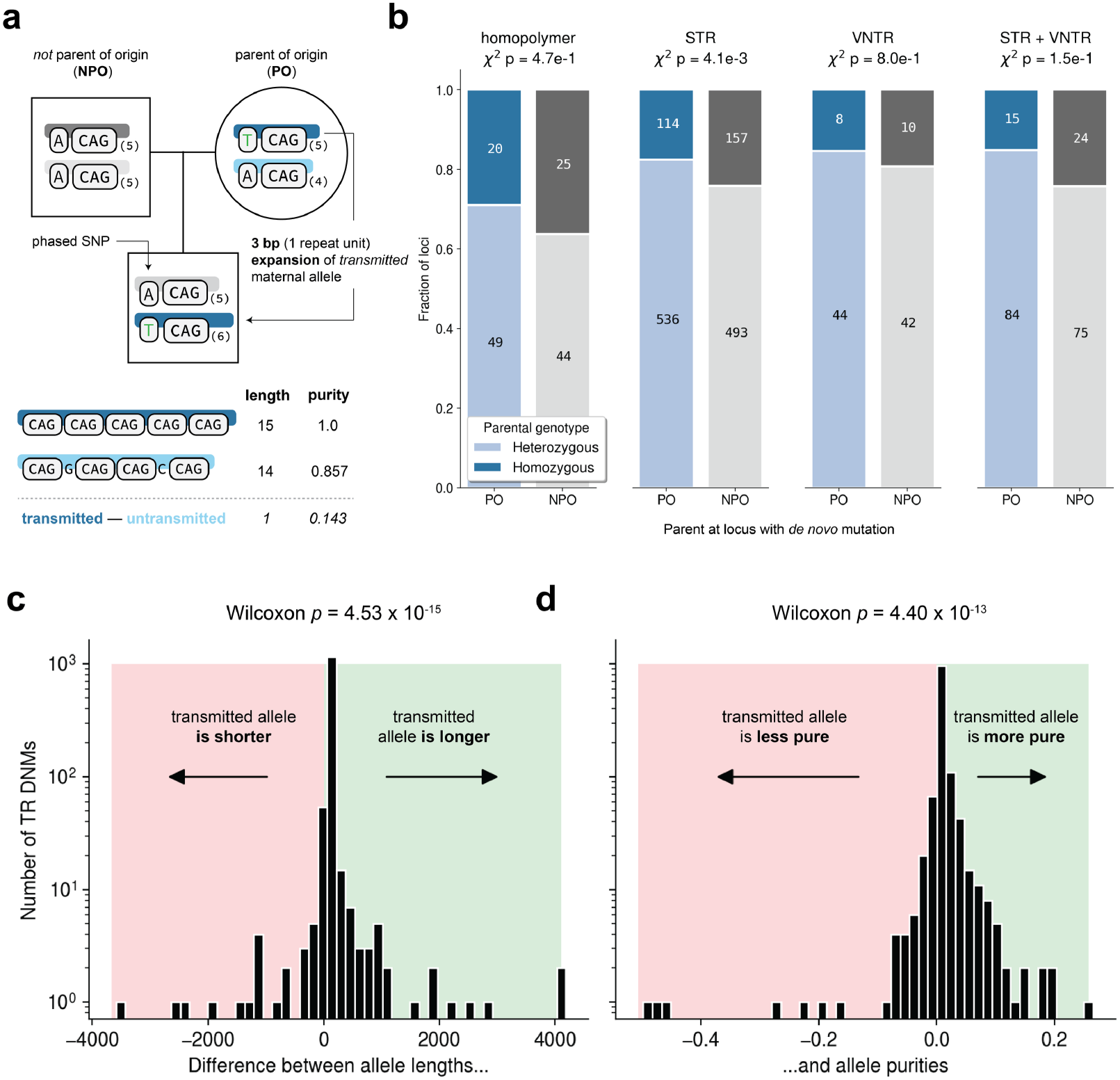
*De novo* mutations are more likely to occur at TR loci that are heterozygous, long, and pure. **a)** At a “toy” tandem repeat locus comprising `CAG` trinucleotide motifs, the child possesses a *de novo* allele with six copies of the `CAG` motif. In this example, we’ve used a flanking single-nucleotide polymorphism to infer that the *de novo* allele was an expansion of a five-copy allele in the maternal germline; thus, the mother is considered the parent-of-origin (PO) and the father is not the parent-of-origin (NPO). By identifying the precursor allele in the PO, we can compare the length and purity of the two tandem repeat alleles in the PO’s genome. Allele length (AL) represents the total number of nucleotides in the allele, and allele purity (AP) represents the fraction of nucleotides in the allele that can be attributed to the expected motif(s). **b)** At a subset of *de novo* TR expansions or contractions for which we could identify the PO, we asked whether the PO and NPO were heterozygous or homozygous for a particular TR allele length at the locus. To test whether DNMs were more likely to occur at heterozygous sites than expected by chance, we compared the fraction of loci at which the PO was heterozygous to the fraction at which the NPO was heterozygous; we used a Chi-square contingency test to determine if the count of DNMs that occurred at heterozygous loci in the PO was significantly higher than expected by chance. We grouped TR loci according to their motif composition, regardless of whether a given locus was simple or complex. Therefore, a “non-homopolymer STR” locus could contain either one or multiple unique motifs, as long as those motifs were all non-homopolymer STRs. Each bar is annotated with the count of mutations in each category. At every tandem repeat locus with a *de novo* mutation, we computed the difference between the **c)** length and **d)** purity of the mutated (transmitted) and untransmitted alleles in the PO. We used a one-tailed Wilcoxon signed-rank test to calculate whether the transmitted alleles were longer and purer, on average, than untransmitted alleles.

### TR mutation rates may be elevated in centromere satellite sequences

The T2T/CHM13 reference assembly contains nearly 200 Mbp of sequence absent from the GRCh38 assembly [50], and almost three quarters of the new sequence comprises centromeric satellites (“CenSat”). Among our tandem repeat *de novo* mutation calls in CHM13, we identified 350 DNMs that overlapped CenSat annotations. Within regions annotated as “centromeric transition” sequences, TR mutation rates were similar to those observed in non-CenSat sequences (**Supp. Fig. 5**). However, TR mutation rates were elevated in other annotated CenSat sequences, namely in rDNA and “Classical human” centromeric satellites (**Supp. Fig. 5**). We also found that STR DNMs were enriched on the *p* arms of the acrocentric chromosomes, likely due to the high density of repetitive satellite sequences in these regions. We validated a small subset of the non-homopolymer STR DNMs overlapping CenSat annotations (*n* = 14) using Element AVITI sequencing data [16]. Compared to a validation rate of 78% (118 of 152) for non-homopolymer STR DNMs outside of CenSat annotations, only 50% (7 of 14) of non-homopolymer STR DNMs within CenSat annotations were validated by Element data. Similarly, we were able to infer a parent-of-origin at only 23% (80 of 350) of TR DNMs within CenSat annotations, compared to nearly 90% (824 of 920) of those outside CenSat annotations (824 of 920). These results suggest that, like *de novo* single-nucleotide mutation rates [16,51,52], TR *de novo* mutation rates may be elevated within centromeric satellite sequences and on the *p* arms of the acrocentric chromosomes. Critically, though, a larger fraction of these TR DNMs are likely false positives, making it challenging to interpret such a mutation rate enrichment without additional data.

### TR mutations occur more frequently in the paternal germline and accumulate with paternal age

Among the 904 TR DNMs for which we inferred a parent-of-origin, 68% were assigned to the father’s germline. In line with prior work [17,38,53], we observed that the number of germline *de novo* STR DNMs (not including pure homopolymer loci) increased with paternal age (effect of 0.68 DNMs per year, *p* = 8 x 10^-3^; **Fig. 4**). As observed in another recent analysis [54], we did not find a significant effect of paternal age on homopolymer DNMs, and we did not observe a maternal age effect on any class of TR DNMs. However, given the relatively low counts of TR DNMs phased to the maternal germline, we may be underpowered to detect such an effect.

**Figure 4:**
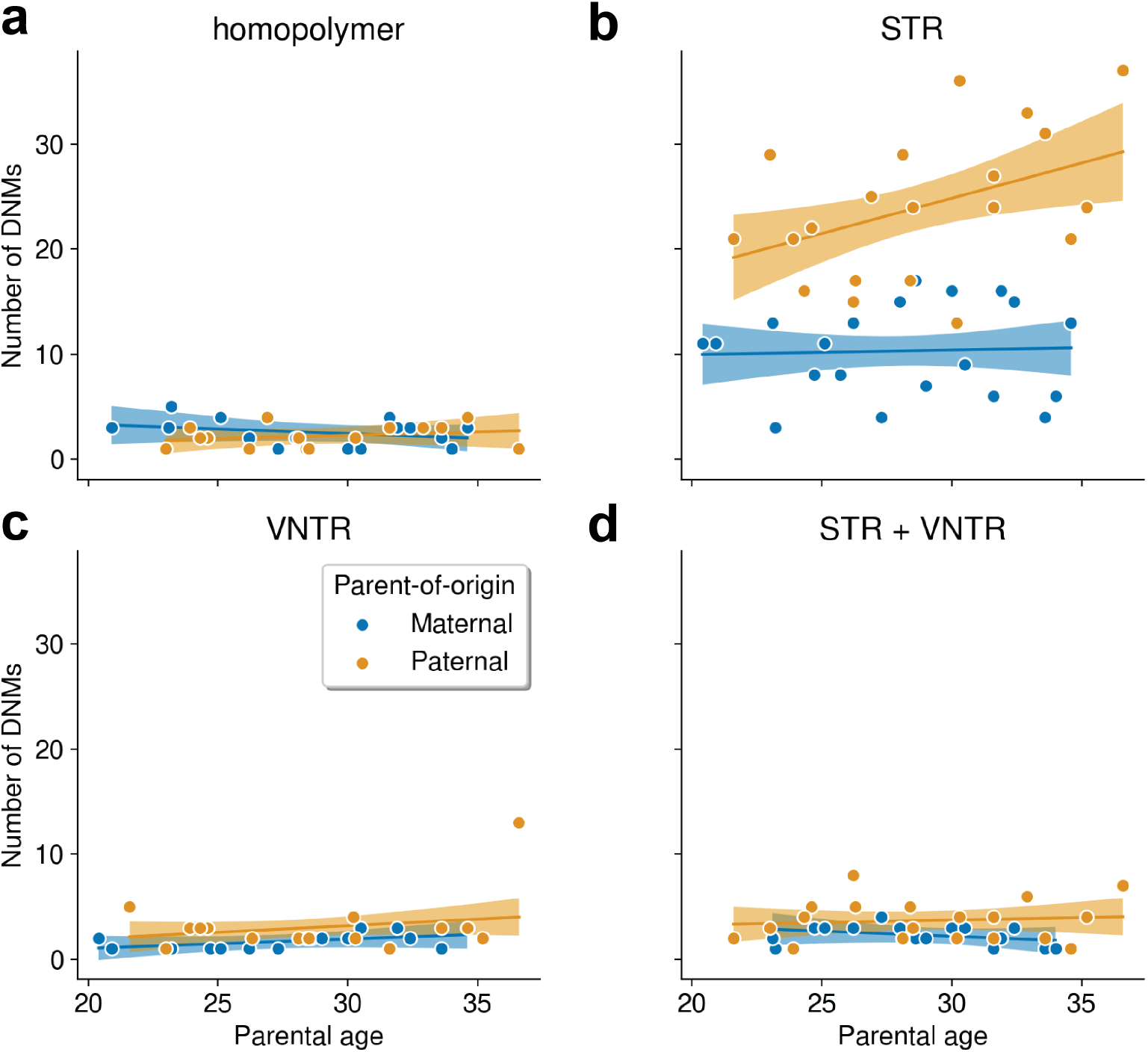
*De novo* short tandem repeat mutations accumulate with paternal age. Correlation between the number of TR DNMs composed of **a)** homopolymer, **b)** STR, **c)** VNTR, or **d)** both STR and VNTR motifs and parental age, stratified by inferred parent-of-origin. Loci in panel **a)** exclusively comprise homopolymer motifs, while loci in panel **b** may contain both homopolymer and non-homopolymer STRs. Loci in panels **b-c** may be simple or complex. Regression lines and 95% confidence intervals are derived from a generalized Poisson linear model using an identity link function.

### Some TR loci are hyper-mutable

The majority of *de novo* TR alleles were private to a single member of the K1463 pedigree, but we previously identified 32 “hyper-mutable” loci that expanded or contracted up to twelve times across the family [16]. Using our updated TR DNM callset, we identified 43 hyper-mutable loci in K1463, of which 16 involved three or more unique *de novo* expansions or contractions. Why were some tandem repeat loci hotspots for mutagenesis, while millions of others remained unchanged from generation to generation? We hypothesized that these hyper-mutable loci contained specific TR motifs that were more prone to expansions and contractions, driving recurrent mutation across the K1463 pedigree. Using TRViz [55], we decomposed the motif structures of all 43 hyper-mutable loci in K1463. Approximately two-thirds of hyper-mutable loci (*n =* 26) were complex, and at 22 of these complex hyper-mutable loci, every *de novo* mutation involved an expansion or contraction of a single motif. For example, at a complex and hypermutable locus near the long non-coding RNA *LINC03021* on chromosome 8, we observed ten unique *de novo* expansions or contractions in K1463. Each of the *de novo* mutations at *LINC03021* involved an expansion or contraction of a specific 19 base pair motif (denoted “a” and illustrated with a blue rectangle in **Fig. 5**). A subtly different version of that motif — comprising just two additional nucleotides, denoted “b” and shown as a light orange square in **Fig. 5** — was stably inherited throughout the pedigree. We also genotyped 100 members of the Human Pangenome Reference Consortium (HPRC) [56] at the *LINC03021* locus and decomposed the motif structure of each individual’s diploid tandem repeat alleles. These HPRC alleles contained more variable counts of the 19bp motif than the similar 21bp motif (**Fig. 5**).

**Figure 5:**
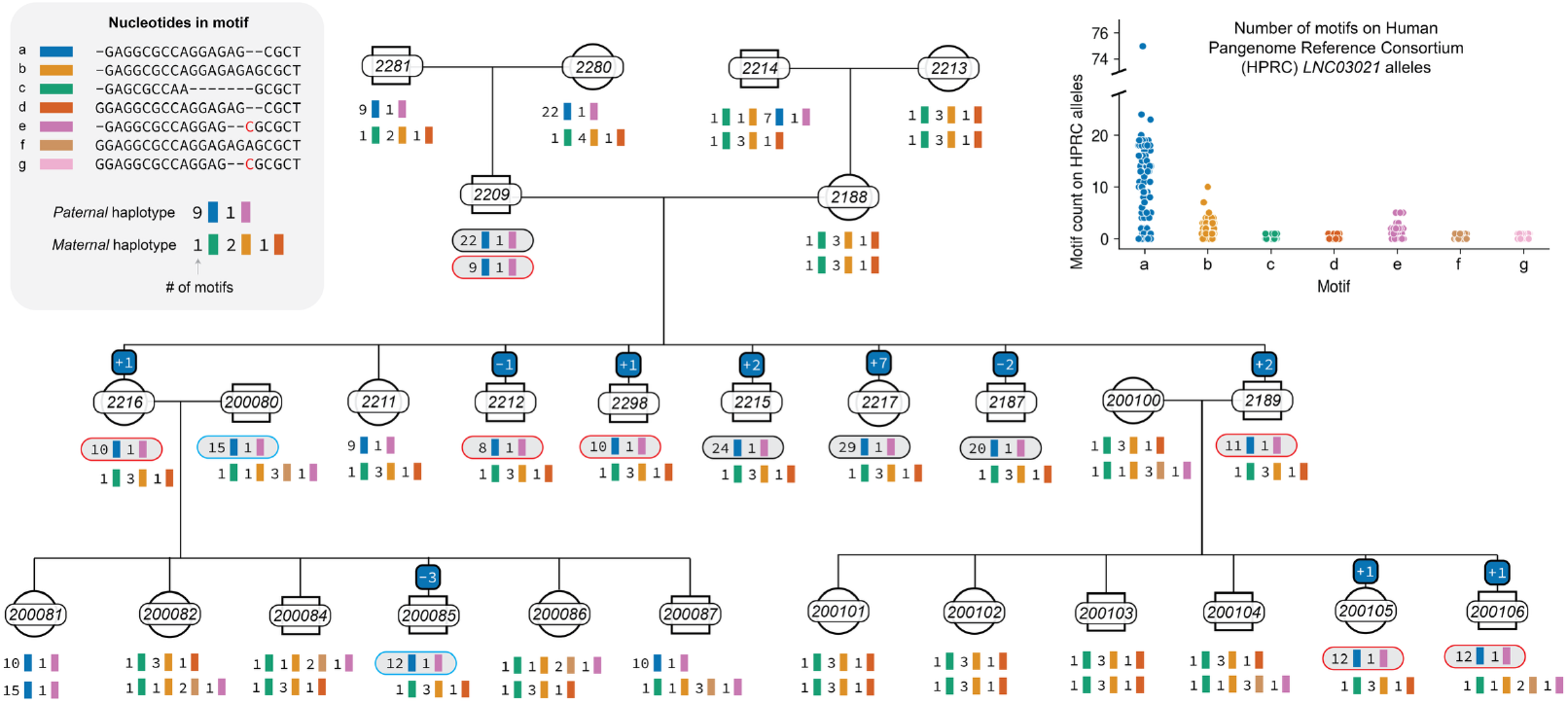
Motif-specific expansions and contractions at a hyper-mutable tandem repeat locus. Tandem repeat alleles are shown for every member of the K1463 pedigree at a single hyper-mutable locus near the long non-coding RNA *LINC03021* (`chr8:2376919-2377075` in T2T/CHM13 coordinates). TR alleles were decomposed into their constituent motifs using TRviz [55], and each motif is shown as a colored square. Motifs that were non-polymorphic within the CEPH 1463 pedigree are not shown. Paternal and maternal haplotypes are shown for each individual in the pedigree. If we observed a *de novo* TR allele in a child, we indicate the number of motifs that expanded or contracted in a small box above the child’s pedigree icon; the box is color-coded according to the specific motif that mutated. In children with *de novo* TR alleles, we also indicate the specific parental haplotype on which the *de novo* expansion or contraction occurred using a gray box with a color-coded outline. For example, the 1-motif expansion observed in sample 2216 occurred on sample 2209’s maternal haplotype. The inset plot illustrates the counts of each motif on a collection of 200 TR alleles (from 100 diploid individuals) in the Human Pangenome Reference Consortium (HPRC). Each point represents the count of a particular motif on an individual TR allele in the HPRC.

While nucleotide content can influence mutability, tandem repeat allele length is also correlated with *de novo* TR mutation rates (**Fig. 3c**). At loci like the one near *LINC03021*, certain motifs may be more mutable simply because they comprise longer uninterrupted arrays on parental TR alleles. To test this hypothesis, we used TRViz to decompose motif structures at 472 complex loci in K1463, including 450 loci with a single DNM and 22 complex hyper-mutable loci; we limited our analysis to complex loci at which we could confidently infer both a parent and parental haplotype-of-origin for every *de novo* allele. At each of these loci, we calculated the relative abundance of each motif on the parental “precursor” allele — that is, the allele that mutated in the parent-of-origin’s germline — and on the untransmitted allele from the same parent (**Supp. Fig. 6**). The motif that mutated to produce a *de novo* allele was the most abundant motif on 474 of 508 precursor alleles, much more often than expected by chance (**Supp. Fig. 6**). This effect was even more pronounced at hyper-mutable loci, where the motif that mutated was the most abundant on 57 of 57 precursor alleles. Motifs that expanded or contracted were also significantly more abundant on precursor alleles than on untransmitted alleles (Wilcoxon signed-rank *p* = 1.2 x 10^-22^; **Supp. Fig. 6**). These results suggest that allele length (and more specifically, the lengths of specific motif arrays) drives mutability at complex TR loci, as well.

## Discussion

Expansions and contractions at TR loci underlie both monogenic and polygenic disease, but germline *de novo* TR mutations have historically been challenging to detect at scale [7]. To comprehensively investigate the factors that influence TR mutagenesis, we sequenced a large, four-generation CEPH/Utah family with the PacBio HiFi long-read platform and genotyped nearly 8 million TR loci using new, highly-sensitive software methods. While TR alleles at most of these loci remained unchanged from generation to generation, we discovered over a thousand germline *de novo* expansions and contractions in the CEPH/Utah K1463 pedigree. TR loci mutated at an average rate of 4.30 x 10^-6^ per locus, per haplotype, per generation (95% CI = 4.1 - 5.0 x 10^-6^), three orders of magnitude higher than the single-nucleotide mutation rate in the same family (**Fig. 1**) [16]. Using long reads, we performed a detailed analysis of the factors that influence TR mutability and discovered a subset of hyper-mutable loci in K1463. Compared to our previous analysis of *de novo* mutations in K1463 [16], we nearly doubled our callset of TR DNMs, discovered additional hyper-mutable loci, and precisely measured the effects of parental genotype, allele length, and allele purity on TR mutability.

Among the 1,270 *de novo* TR mutations we observed in K1463, 1,078 exclusively involved short tandem repeat (STR) motifs. We observed similar numbers of expansions and contractions overall, though we found an enrichment of contractions over expansions at loci comprising dinucleotide motifs (**Fig. 2**). Many prior studies have instead documented a bias toward *expansions* at STR loci [17,18], and we hypothesized that the composition of our TR catalog (which contained more and larger loci) might explain the observed enrichment of contractions. After genotyping a separate catalog of STR loci — which comprised shorter TR loci, on average — with our PacBio HiFi sequencing reads, we still observed similar numbers of expansions and contractions at most loci, and significantly more contractions than expansions at dinucleotide loci. As a result, we cannot attribute the lack of an observed expansion bias to differences in TR catalog composition alone. Instead, we hypothesize that differences in our sequencing and genotyping strategies (and our use of long PacBio sequencing reads, in particular) are likely responsible for the discordance between ours and previous results.

We found that *de novo* TR mutations were more likely to occur in the germline of a heterozygous parent (**Fig. 3**). Because heterozygosity at a TR locus (that is, the presence of more than one unique TR allele) is the outcome of at least one prior *de novo* mutation event, we further hypothesized that TR loci with *de novo* mutations in K1463 would also exhibit higher population-level heterozygosity in an unrelated cohort of human genomes. Supporting this hypothesis, we found that *de novo* expansions and contractions in K1463 occurred at TR loci that were significantly more heterozygous in the Human Pangenome Reference Consortium (HPRC) than loci that did not mutate (**Supp. Fig. 4**). These results lend support to the “heterozygote instability” hypothesis, which posits that *de novo* mutations are more likely to occur when there is a large difference in TR allele lengths between homologous chromosomes [17,57–59].

By phasing TR mutations with long-read HiFi sequencing data, we identified the parental alleles that mutated to produce a *de novo* expansion or contraction in each child. These “precursor” alleles were longer and purer, on average, than the parental TR alleles that were *not* transmitted to a subsequent generation (**Fig. 3c-d**). This observation is in line with evidence that interrupted TR alleles, in which the repetitive motif array is interrupted by single-nucleotide substitutions or small insertions and deletions, are less likely to undergo slipped-strand mispairing [27,30]. We also found that older fathers contribute more STR DNMs than younger fathers (**Fig. 4**), consistent with previous findings using short reads [38,54]. We did not observe a significant paternal age effect on expansions or contractions at homopolymers, and possibly owing to the relatively small number of *de novo* mutations at variable number of tandem repeats (VNTRs), we did not find a parental age effect on VNTR mutation. By sequencing additional families with long-read platforms, we may be able to improve our estimates of parental age effects on *de novo* mutation rates at homopolymer and VNTR loci.

In our updated TR DNM callset, we discovered 43 TR loci at which we observed two or more *de novo* expansions or contractions in K1463. Given the relatively small numbers of these “recurrent” DNMs in K1463, it is difficult to identify specific features of TR loci that make them prone to hyper-mutation. At a complex locus with 7 unique motifs near the long non-coding RNA *LINC03021* (T2T/CHM13 coordinates chr8:2623352-2623487), we discovered that *de novo* expansions and contractions exclusively involved a particular 19-base pair motif. A subtly different version of that motif — differing by just 2 base pairs — was stably inherited from generation to generation and was less variable on HPRC alleles at the same locus (**Fig. 5)**. This result suggested that motif-specific mutability might be explained by small differences in nucleotide content; however, we also considered the possibility that motifs might mutate more often if they comprise longer arrays of uninterrupted sequence. To test this second hypothesis, we decomposed the motif structures of TR alleles at hundreds of complex loci in K1463. At these complex loci, the motifs that expanded or contracted were almost always more abundant than stably inherited motifs on parental “precursor” alleles (**Supp. Fig. 6**). Thus, the mutability of specific motifs at recurrent DNMs could potentially be explained by the fact that they comprise longer, uninterrupted arrays of repetitive nucleotide content. Future work will be required to disentangle the effects of nucleotide content and allele length on the relative mutability of TR motifs.

In our original analysis of TR mutation in K1463, we observed unusually high numbers of *de novo* TR alleles in generation 4 (G4). The G4 DNMs were also enriched for short expansions and contractions at homopolymers and exhibited a striking parent-of-origin bias (see **Supplementary Note** for more details). Using a series of downsampling experiments, we demonstrated that our genotyping and *de novo* mutation detection pipeline was highly sensitive to allelic dropout. These challenges were mitigated once the parents of G4 were re-sequenced to ∼70X depth. Our results suggest that accurate genotyping and parent-of-origin inference at *de novo* TR alleles require very high parental long-read sequencing depth (on the order of 60-70X) to avoid allelic dropout. Short-read sequencing platforms likely suffer from dropout at heterozygous loci where one TR allele is much longer than the other and exceeds the length of a typical paired-end read. By performing similar downsampling experiments using short-read datasets, we may be able to determine whether allelic dropout is a pervasive problem for TR genotyping and *de novo* mutation detection.

As we continue to sequence large, multi-generational families with new technologies and develop improved software methods for *de novo* TR detection, we will be able to more precisely resolve the dynamics of mutagenesis at TRs. TRs comprise highly repetitive DNA sequences that confound sequence alignment, and short reads are unable to completely span TR loci that measure hundreds or thousands of base pairs. Even relatively small loci may undergo large expansions, giving rise to new alleles that exceed the length of short reads. Long read sequencing technologies, such as the Pacific Biosciences (PacBio) and Oxford Nanopore Technologies (ONT) platforms, generate reads that can completely capture both reference and mutated tandem repeat alleles. Long reads can also be more reliably mapped to telomeres, centromeres, and other low-complexity sequences, enabling comprehensive genotyping of TR variation throughout the genome [60]. New short-read technologies, like the Element AVITI platform, exhibit very low error rates at homopolymers [54,61], making them well-suited for genotyping expansions and contractions at homopolymer loci. In this study, our long-read sequencing approach enabled comprehensive genotyping of large TRs, including VNTRs and “complex” loci that contain both STRs and VNTRs. Using long reads, we also precisely identified the parental allele on which a majority of *de novo* TR mutations occurred, enabling us to directly measure the relationship between allele purity and length on TR mutability. About half of the TR DNMs in K1463 produced *de novo* alleles larger than a typical short read length and would have been impossible to find using short-read platforms. By applying long-read technologies to other pedigrees, we expect to uncover even more hyper-mutable TRs, making it possible to identify the genomic factors that make these loci such potent engines of mutagenesis.

## Materials and Methods

### Additional PacBio HiFi sequencing and alignment

As described in the **Supplementary Note**, we performed “top-up” sequencing to generate additional HiFi read coverage for four members of CEPH K1463. Top-up sequencing data was generated for samples NA12879, NA12886, 200080, and 200100. DNA was extracted from whole blood using the Flexigene system (Qiagen 51206) and whole-genome sequencing was performed with the PacBio HiFi platform as previously described [16].

We aligned the unmapped BAM files generated by each PacBio HiFi run to both the CHM13v2 or GRCh38 reference assemblies using `pbmm2` v1.17.0:

~~~
/path/to/pbmm2 align --num-threads $threads --sort --sort-memory $mem
--preset CCS --sample $sample_name --bam-index BAI --unmapped $REF
$INPUT_BAM $OUTPUT_BAM
~~~

### Tandem repeat genotyping

We used the Tandem Repeat Genotyper (TRGT) v3.0.0 [43] to genotype each member of the CEPH K1463 at a catalog of approximately 7.8 million unique tandem repeat loci with the following command:

~~~
/path/to/trgt genotype --threads $threads --genome $REF --repeats $BED
--reads $BAM --output-prefix $prefix --karyotype $karyotype
~~~

The $karyotype flag was set to “XX” for female samples and “XY” for male samples.

Methods used to create our catalog of TR loci are described in a previous manuscript [16]. Briefly, we used Tandem Repeats Finder [47] and TRsolve to identify all tandem repeat sequences measuring between 10 and 10,000bp in the GRCh38 and T2T CHM13v2 reference assemblies. Any TR loci within 50 base pairs of each other were merged into a single, “complex” TR locus.

### Single-nucleotide variant calling

We used germline single-nucleotide variants to perform haplotype phasing and infer the parent and/or haplotype-of-origin for *de novo* tandem repeat alleles. We used Singularity v4.1.1 [62] and DeepVariant v.1.9.0 [45] to genotype SNVs in each sample using the following command:

~~~
singularity exec –-cleanenv -H $TMPDIR -B
/usr/lib/locale/:/usr/lib/locale/ $SIF run_deepvariant --model_type
PACBIO --num_shards $shards --output_vcf $OUTPUT_VCF --output_gvcf
$OUTPUT_GVCF --reads $BAM --ref $REF --regions $CHROM --sample_name
$sample_name --make-examples-extra-args “select_variant_types=‘snps’,min_mapping_quality=1
~~~

### Haplotype phasing

We jointly phased SNV and TR genotypes in each sample using HiPhase v1.4.4 [46] using the following command:

~~~
/path/to/hiphase --threads $threads --bam $INPUT_BAM --vcf $SNV_VCF --
vcf $STR_VCF --reference $REF --output-vcf $PHASED_SNV_VCF --output-
vcf $PHASED_STR_VCF
~~~

### Identifying candidate de novo TR expansions and contractions

We used TRGT-denovo v0.2.3 [44] to identify candidate *de novo* tandem repeat alleles in each child in G3 and G4 using the following command:

~~~
/path/to/trgt-denovo trio –reference $REF –bed $REPEAT_CATALOG –father
$DAD_PREFIX –mother $MOM_PREFIX –child $KID_PREFIX –out $OUTPUT -@
$THREADS
~~~

As described in our previous manuscript [16], we applied a set of basic filters to eliminate likely false positive *de novo* mutation calls:

- Spanning sequencing depth in the child and both parents >= 10 reads
- At least 2 reads of support for the *de novo* allele, and at least 20% of all reads supporting the *de novo* allele
- A maximum of 5% of total parental read support for the *de novo* allele

We also removed candidate DNMs observed in G3 if the *de novo* allele length was observed in a grandparent.

### Inferring the parent- and haplotype-of-origin for de novo TR mutations

We inferred the likely parent-of-origin for each *de novo* TR allele using the same approach described in [16]. Briefly, we identified “informative” single-nucleotide variants (SNVs) within 500 kbp upstream or downstream of the *de novo* allele. These informative sites were SNVs at which the child was genotyped as heterozygous and the parental genotypes did not match (e.g., child’s genotype is `0/1`, mother’s genotype is either `0/1` or `1/1`, and father’s genotype is `0/0`.) In this example, we can infer that the alternate allele in the child was inherited from the mother and the reference allele was inherited from the father. At each of these informative sites, we then examined the jointly phased STR and SNV genotypes in the child. For all informative heterozygous sites in the child that were assigned the same HiPhase phase block as the phased STR locus in the child, we asked whether the reference or alternate SNV allele was present on the same haplotype as the *de novo* TR allele. In other words, at the example informative site above, if the child’s phased SNV genotype was `1|0`, their phased genotype at the nearby TR locus was `2|1`, and the *de novo* TR allele was the `2` allele, we would infer that the *de novo* mutation occurred in the mother.

To infer the parental haplotype of origin, we could then examine the phased SNV genotypes in the mother at this example informative site. If the mother’s phased SNV genotype was `0|1`, then we can infer that the *de novo* mutation occurred on mom’s “second” haplotype (the order of haplotypes being dependent on HiPhase output, and not carrying any explicit biological meaning). Assuming the mother’s phased TR genotype was `1|0` and the phased TR genotype was part of the same phase block as the mother’s phased SNV genotype, we can further infer that the `0` TR allele in the mother was the allele that mutated.

### Determining the size of de novo expansions and contractions at tandem repeats

We used two methods to determine the likely size of every *de novo* TR allele. The first, which we refer to as the “parsimony-based” approach, simply involves comparing the size of the *de novo* allele to the sizes of the parental alleles at the same locus. If we are able to determine a confident parent-of-origin for the *de novo* mutation, we compare the size of the *de novo* allele to the sizes of the diploid alleles in that parent-of-origin. If we cannot determine a parent-of-origin, we compare the size of the *de novo* allele to the sizes of the diploid alleles in both parents. We calculate the difference in size between the *de novo* allele and each of the parental alleles, and assume that the smallest absolute difference is the most parsimonious *de novo* mutation size. For example, given an autosomal *de novo* allele that measures 20 base pairs in a child, and assuming the father’s TR alleles measure 25 and 40 base pairs and the mother’s TR alleles measure 30 and 35 base pairs, we would infer the most parsimonious change to be a 5bp contraction in the father’s germline. The second method, which we refer to as the “precursor-based” method, is only applicable when we have inferred both the parent-of-origin *and* haplotype-of-origin for a *de novo* TR mutation. Using this method, we directly compare the length of the *de novo* allele in the child to the length of the TR allele on the parental haplotype on which the *de novo* mutation was inferred to occur.

### Identifying recurrently mutated TR loci

As reported previously [16], we searched for recurrent TR DNMs by first curating a list of all TR loci at which TRGT-denovo reported at least two *de novo* alleles in G3 and/or G4. All *de novo* alleles were required to pass the TR DNM filters listed above (see section “Identifying candidate *de novo* TR expansions and contractions”). At the loci with recurrent DNMs, we further required that every member of the CEPH K1463 pedigree was genotyped by TRGT with at least 10 HiFi reads supporting the genotype assignment (i.e., the sum of the `SD` FORMAT field must be greater or equal to 10). These filtering criteria produced a candidate list of 78 recurrent DNMs. We then manually inspected the HiFi read evidence supporting each recurrent *de novo* TR allele to remove potential false positives.

### Decomposing tandem repeat alleles for visualization

Our tandem repeat catalog comprises approximately 7.8 million unique tandem repeat loci in the T2T/CHM13v2 reference genome sequence. We used Tandem Repeats Finder (TRF) or TRsolve to determine the motif composition of each locus. For example, a homopolymer locus might have a motif structure like (A)_n_, where *n* indicates a variable number of motifs; a complex locus might have a motif structure like (AG)_n_(CCCGCGG)_n_. After genotyping a tandem repeat locus in a diploid genome with TRGT [43], we know both the nucleotide sequences and lengths of both TR alleles. However, we do not know the motif composition of each allele. In order to “decompose” an allele’s nucleotide sequence into its most likely motif composition, we used the TRViz software tool [55]. TRViz accepts a tandem repeat allele sequence as input, along with a list of the expected motifs that comprise the allele; the former was derived from TRGT output, and the latter from the motifs in our TR catalog. TRViz then determines the most likely structure of the allele sequence, using both the “reference” motifs as well as *de novo* motifs that might produce a more accurate motif decomposition. For example, if the allele sequence is TTTTAGAGAGAC, and the expected motifs at the locus were `T` and `AG`, TRViz might infer the presence of a *de novo* `AC` motif. This approach was used for visualizing the motif structure of tandem repeat alleles at hyper-mutable loci (see **Fig. 5**). To identify the specific motifs that mutated to produce a *de novo* allele, we applied a second approach (described below) that involved making a direct comparison between a *de novo* allele and its “precursor” allele in the parent-of-origin.

### Identifying the motifs that mutated within complex TR loci

Complex tandem repeat (TR) loci are regions comprising multiple distinct TR motifs that are proximal, overlapping, or nested. When TRGT-denovo identifies DNMs within such regions, it reports the overall length change for the locus. To pinpoint the specific TR motif that mutated within a complex locus, we first decomposed each complex TR locus to determine the coordinate ranges of the individual TR annotations in the parental allele. Motif decomposition was performed using the repeat identification tool “ribbit” [63], which reports the coordinates of individual TR annotations along with their respective motifs, purity scores, and sub-coordinate ranges of perfect repeat stretches. The *de novo* allele is then aligned to the parental allele to identify sequence variations (indels). Each sequence variation is assigned to a TR motif based on the parental motif annotated at that coordinate. Deletions were directly assigned based on overlap with annotated motifs. For insertions occurring at the junction between two motifs, the inserted sequence was aligned to the perfect repeat stretches of both candidate TR motifs. The insertion was assigned to the TR motif with the highest number of matching bases.

## Data availability

Sequencing data for the 28 members of the K1463 pedigree are available as part of the AWS Open Data Program, the European Nucleotide Archive (ENA), or the Database of Genotypes and Phenotypes (dbGaP). Aligned BAM files for 23 members of K1463 (who consented to broad research access) are available in the AWS Open Data Program (s3://platinum-pedigree-data/) as well as the European Nucleotide Archive (BioProject: PRJEB86317). Aligned BAM files for the remaining five members of the pedigree (who did not consent to public access) are available in dbGaP (phs003793.v1.p1; Platinum Pedigree Consortium LRS). Aligned BAM files containing “top-up” sequencing data for four members of K1463 (who consented to broad research access) will be uploaded to both the AWS Open Data Program and ENA, using the accessions provided above.

## Code availability

We have uploaded *snakemake* [64] workflows for alignment, variant calling, TR discovery, phasing, and all analyses presented in this manuscript to a GitHub repository (URL: https://github.com/tomsasani/ceph-k1463-tandem-repeats).

## Acknowledgements

We thank the members of CEPH/Utah family K1463, who supported this analysis (among many others) through their research participation. We also thank Drs. Raymond White, Jean-Marc Lalouel, and Mark Leppert, who were critically important for the original engagement of the CEPH/Utah families. Research reported in this publication was supported, in part, by the National Human Genome Research Institute (NHGRI) of the National Institutes of Health (NIH) under grants 5R00HG012796-04 (to H.D. and A.A.), R35GM118335 (to L.B.J.); and R01HG002385 and R01HG010169 (to E.E.E.). H.D. and A.A. are also supported by NHMRC Investigator grant GNT2026126. The computational resources used at the University of Utah were partially funded by the NIH Shared Instrumentation Grant 1S10OD021644-01A1. The content is solely the responsibility of the authors and does not necessarily represent the official views of the NIH. E.E.E. is an investigator of the Howard Hughes Medical Institute.

## Author contributions

Contributions are described following the Contributor Role Taxonomy (CRediT) framework: https://credit.niso.org/

TAS: Methodology, Software, Formal Analysis, Writing - Original Draft.

MEG: Conceptualization, Formal Analysis, Writing - Reviewing & Editing.

AKA: Methodology, Software.

TJN: Formal Analysis, Writing – Reviewing & Editing.

DWN: Resources, Project Administration.

ED: Methodology, Software.

TM: Methodology, Software.

KMM: Resources.

KH: Resources.

MA: Resources.

EJK: Methodology, Software.

DP: Methodology.

ZK: Resources.

GSB: Resources, Methodology.

WJR: Resources.

LBJ: Resources.

CEM: Resources, Writing – Reviewing & Editing.

MAE: Resources.

PNV: Methodology, Writing - Reviewing & Editing.

EEE: Conceptualization, Writing - Reviewing & Editing.

ARQ: Conceptualization, Writing - Reviewing & Editing, Supervision.

HD: Conceptualization, Methodology, Writing - Review & Editing, Supervision.

## References

1. Hiatt L, Weisburd B, Dolzhenko E, Rubinetti V, Avvaru AK, VanNoy GE, et al. STRchive: a dynamic resource detailing population-level and locus-specific insights at tandem repeat disease loci. Genome Med. 2025;17: 29.

2. Bates GP. History of genetic disease: the molecular genetics of Huntington disease - a history. Nat Rev Genet. 2005;6: 766–773.

3. Ishiura H, Doi K, Mitsui J, Yoshimura J, Matsukawa MK, Fujiyama A, et al. Expansions of intronic TTTCA and TTTTA repeats in benign adult familial myoclonic epilepsy. Nat Genet. 2018;50: 581–590.

4. Gall-Duncan T, Sato N, Yuen RKC, Pearson CE. Advancing genomic technologies and clinical awareness accelerates discovery of disease-associated tandem repeat sequences. Genome Res. 2022;32: 1–27.

5. Beyter D, Ingimundardottir H, Oddsson A, Eggertsson HP, Bjornsson E, Jonsson H, et al. Long-read sequencing of 3,622 Icelanders provides insight into the role of structural variants in human diseases and other traits. Nat Genet. 2021;53: 779–786.

6. Mukamel RE, Handsaker RE, Sherman MA, Barton AR, Zheng Y, McCarroll SA, et al. Protein-coding repeat polymorphisms strongly shape diverse human phenotypes. Science. 2021;373: 1499–1505.

7. Tanudisastro HA, Deveson IW, Dashnow H, MacArthur DG. Sequencing and characterizing short tandem repeats in the human genome. Nat Rev Genet. 2024;25: 460–475.

8. Levinson G, Gutman GA. Slipped-strand mispairing: a major mechanism for DNA sequence evolution. Mol Biol Evol. 1987;4: 203–221.

9. Fan H, Chu J-Y. A brief review of short tandem repeat mutation. Genomics Proteomics Bioinformatics. 2007;5: 7–14.

10. Verbiest M, Maksimov M, Jin Y, Anisimova M, Gymrek M, Bilgin Sonay T. Mutation and selection processes regulating short tandem repeats give rise to genetic and phenotypic diversity across species. J Evol Biol. 2023;36: 321–336.

11. Jakupciak JP, Wells RD. Genetic instabilities in (CTG.CAG) repeats occur by recombination. J Biol Chem. 1999;274: 23468–23479.

12. Richard GF, Pâques F. Mini- and microsatellite expansions: the recombination connection. EMBO Rep. 2000;1: 122–126.

13. Huang Q-Y, Xu F-H, Shen H, Deng H-Y, Liu Y-J, Liu Y-Z, et al. Mutation patterns at dinucleotide microsatellite loci in humans. Am J Hum Genet. 2002;70: 625–634.

14. Debrauwère H, Buard J, Tessier J, Aubert D, Vergnaud G, Nicolas A. Meiotic instability of human minisatellite CEB1 in yeast requires DNA double-strand breaks. Nat Genet. 1999;23: 367–371.

15. Jeffreys AJ, Neil DL, Neumann R. Repeat instability at human minisatellites arising from meiotic recombination. EMBO J. 1998;17: 4147–4157.

16. Porubsky D, Dashnow H, Sasani TA, Logsdon GA, Hallast P, Noyes MD, et al. Human de novo mutation rates from a four-generation pedigree reference. Nature. 2025;643: 427– 436.

17. Mitra I, Huang B, Mousavi N, Ma N, Lamkin M, Yanicky R, et al. Patterns of de novo tandem repeat mutations and their role in autism. Nature. 2021;589: 246–250.

18. Steely CJ, Watkins WS, Baird L, Jorde LB. The mutational dynamics of short tandem repeats in large, multigenerational families. Genome Biol. 2022;23: 253.

19. Kristmundsdottir S, Jonsson H, Hardarson MT, Palsson G, Beyter D, Eggertsson HP, et al. Sequence variants affecting the genome-wide rate of germline microsatellite mutations. Nat Commun. 2023;14: 3855.

20. Sun JX, Helgason A, Masson G, Ebenesersdóttir SS, Li H, Mallick S, et al. A direct characterization of human mutation based on microsatellites. Nat Genet. 2012;44: 1161– 1165.

21. Sasani TA, Pedersen BS, Gao Z, Baird L, Przeworski M, Jorde LB, et al. Large, three-generation human families reveal post-zygotic mosaicism and variability in germline mutation accumulation. Elife. 2019;8. doi:10.7554/eLife.46922

22. Jónsson H, Sulem P, Kehr B, Kristmundsdottir S, Zink F, Hjartarson E, et al. Parental influence on human germline de novo mutations in 1,548 trios from Iceland. Nature. 2017;549: 519–522.

23. Eslami Rasekh M, Hernández Y, Drinan SD, Fuxman Bass JI, Benson G. Genome-wide characterization of human minisatellite VNTRs: population-specific alleles and gene expression differences. Nucleic Acids Res. 2021;49: 4308–4324.

24. Denoeud F, Vergnaud G, Benson G. Predicting human minisatellite polymorphism. Genome Res. 2003;13: 856–867.

25. Chakraborty R, Kimmel M, Stivers DN, Davison LJ, Deka R. Relative mutation rates at di-, tri-, and tetranucleotide microsatellite loci. Proc Natl Acad Sci U S A. 1997;94: 1041–1046.

26. Brinkmann B, Klintschar M, Neuhuber F, Hühne J, Rolf B. Mutation rate in human microsatellites: influence of the structure and length of the tandem repeat. Am J Hum Genet. 1998;62: 1408–1415.

27. Gacy AM, Goellner G, Juranić N, Macura S, McMurray CT. Trinucleotide repeats that expand in human disease form hairpin structures in vitro. Cell. 1995;81: 533–540.

28. McGinty RJ, Balick DJ, Mirkin SM, Sunyaev SR. Inherent instability of simple DNA repeats shapes an evolutionarily stable distribution of repeat lengths. Nat Commun. 2025;17: 93.

29. Rajan-Babu I-S, Dolzhenko E, Eberle MA, Friedman JM. Sequence composition changes in short tandem repeats: heterogeneity, detection, mechanisms and clinical implications. Nat Rev Genet. 2024;25: 476–499.

30. Goldberg ME, Dashnow H, Harris K, Quinlan AR. The selective dynamics of interruptions at short tandem repeats. bioRxiv. 2025. p. 2025.06.09.658724. doi:10.1101/2025.06.09.658724

31. Payseur BA, Jing P, Haasl RJ. A genomic portrait of human microsatellite variation. Mol Biol Evol. 2011;28: 303–312.

32. Bacolla A, Larson JE, Collins JR, Li J, Milosavljevic A, Stenson PD, et al. Abundance and length of simple repeats in vertebrate genomes are determined by their structural properties. Genome Res. 2008;18: 1545–1553.

33. Kelkar YD, Tyekucheva S, Chiaromonte F, Makova KD. The genome-wide determinants of human and chimpanzee microsatellite evolution. Genome Res. 2008;18: 30–38.

34. Lin Y, Dion V, Wilson JH. Transcription promotes contraction of CAG repeat tracts in human cells. Nat Struct Mol Biol. 2006;13: 179–180.

35. Bowater RP, Jaworski A, Larson JE, Parniewski P, Wells RD. Transcription increases the deletion frequency of long CTG.CAG triplet repeats from plasmids in Escherichia coli. Nucleic Acids Res. 1997;25: 2861–2868.

36. Nakamori M, Pearson CE, Thornton CA. Bidirectional transcription stimulates expansion and contraction of expanded (CTG)*(CAG) repeats. Hum Mol Genet. 2011;20: 580–588.

37. Kayser M, Roewer L, Hedman M, Henke L, Henke J, Brauer S, et al. Characteristics and frequency of germline mutations at microsatellite loci from the human Y chromosome, as revealed by direct observation in father/son pairs. Am J Hum Genet. 2000;66: 1580–1588.

38. Goldberg ME, Noyes MD, Eichler EE, Quinlan AR, Harris K. Effects of parental age and polymer composition on short tandem repeat de novo mutation rates. Genetics. 2024;226. doi:10.1093/genetics/iyae013

39. Legendre M, Pochet N, Pak T, Verstrepen KJ. Sequence-based estimation of minisatellite and microsatellite repeat variability. Genome Res. 2007;17: 1787–1796.

40. Andreassen R, Egeland T, Olaisen B. Mutation rate in the hypervariable VNTR g3 (D7S22) is affected by allele length and a flanking DNA sequence polymorphism near the repeat array. Am J Hum Genet. 1996;59: 360–367.

41. Vergnaud G, Mariat D, Apiou F, Aurias A, Lathrop M, Lauthier V. The use of synthetic tandem repeats to isolate new VNTR loci: cloning of a human hypermutable sequence. Genomics. 1991;11: 135–144.

42. Kronenberg Z, Nolan C, Mokveld T, Rowell WJ, Lee S, Dolzhenko E, et al. The Platinum Pedigree: A long-read benchmark for genetic variants. bioRxiv. 2024. p. 2024.10.02.616333. doi:10.1101/2024.10.02.616333

43. Dolzhenko E, English A, Dashnow H, De Sena Brandine G, Mokveld T, Rowell WJ, et al. Characterization and visualization of tandem repeats at genome scale. Nat Biotechnol. 2024;42: 1606–1614.

44. Mokveld T, Dolzhenko E, Dashnow H, Nicholas TJ, Sasani T, van der Sanden B, et al. TRGT-denovo: accurate detection of de novo tandem repeat mutations. bioRxivorg. 2024. doi:10.1101/2024.07.16.600745

45. Poplin R, Chang P-C, Alexander D, Schwartz S, Colthurst T, Ku A, et al. A universal SNP and small-indel variant caller using deep neural networks. Nat Biotechnol. 2018;36: 983– 987.

46. Holt JM, Saunders CT, Rowell WJ, Kronenberg Z, Wenger AM, Eberle M. HiPhase: jointly phasing small, structural, and tandem repeat variants from HiFi sequencing. Bioinformatics. 2024;40. doi:10.1093/bioinformatics/btae042

47. Benson G. Tandem repeats finder: a program to analyze DNA sequences. Nucleic Acids Res. 1999;27: 573–580.

48. Mousavi N, Shleizer-Burko S, Yanicky R, Gymrek M. Profiling the genome-wide landscape of tandem repeat expansions. Nucleic Acids Res. 2019;47: e90.

49. Liao W-W, Asri M, Ebler J, Doerr D, Haukness M, Hickey G, et al. A Draft Human Pangenome Reference. bioRxiv. 2022. doi:10.1101/2022.07.09.499321

50. Nurk S, Koren S, Rhie A, Rautiainen M, Bzikadze AV, Mikheenko A, et al. The complete sequence of a human genome. Science. 2022;376: 44–53.

51. Noyes MD, Sui Y, Kwon Y, Koundinya N, Wong I, Munson KM, et al. Long-read sequencing of trios reveals increased germline and postzygotic mutation rates in repetitive DNA. bioRxivorg. 2025. doi:10.1101/2025.07.18.665621

52. Lin J, Mastrorosa FK, Noyes MD, Yoo D, Rhie A, Porubsky D, et al. Human acrocentric chromosome short arm de novo mutation and recombination. bioRxivorg. bioRxiv; 2025. p. 2025.12.16.694519. doi:10.64898/2025.12.16.694519

53. Forster P, Hohoff C, Dunkelmann B, Schürenkamp M, Pfeiffer H, Neuhuber F, et al. Elevated germline mutation rate in teenage fathers. Proc Biol Sci. 2015;282: 20142898.

54. Happ HC, Sasani TA, Warner D, Neklason DW, Quinlan AR. AVITI sequencing of a four-generation CEPH/Utah pedigree confirms low mutation rates at homopolymer loci despite their low sequence complexity. bioRxiv. 2025. doi:10.1101/2025.09.25.678675

55. Park J, Kaufman E, Valdmanis PN, Bafna V. TRviz: a Python library for decomposing and visualizing tandem repeat sequences. Bioinform Adv. 2023;3: vbad058.

56. Liao W-W, Asri M, Ebler J, Doerr D, Haukness M, Hickey G, et al. A draft human pangenome reference. Nature. 2023;617: 312–324.

57. Amos W, Sawcer SJ, Feakes RW, Rubinsztein DC. Microsatellites show mutational bias and heterozygote instability. Nat Genet. 1996;13: 390–391.

58. Masters BS, Johnson LS, Johnson BGP, Brubaker JL, Sakaluk SK, Thompson CF. Evidence for heterozygote instability in microsatellite loci in house wrens. Biol Lett. 2011;7: 127–130.

59. Amos W, Kosanović D, Eriksson A. Inter-allelic interactions play a major role in microsatellite evolution. Proc Biol Sci. 2015;282: 20152125.

60. Grigorev K, Foox J, Bezdan D, Butler D, Luxton JJ, Reed J, et al. Haplotype diversity and sequence heterogeneity of human telomeres. Genome Res. 2021;31: 1269–1279.

61. Arslan S, Garcia FJ, Guo M, Kellinger MW, Kruglyak S, LeVieux JA, et al. Sequencing by avidity enables high accuracy with low reagent consumption. Nat Biotechnol. 2024;42: 132–138.

62. Trudgian D, Kurtzer GM cclerget, Bauer M, Kaneshiro I, Godlove D, et al. sylabs/singularity: SingularityCE 4.3.2. Zenodo; 2025. doi:10.5281/ZENODO.5570766

63. Avvaru AK, Sharma A, Sowpati DT. Ribbit: Accurate identification and annotation of complex tandem repeat sequences in genomes. bioRxiv. bioRxiv; 2025. p. 2025.02.06.636828. doi:10.1101/2025.02.06.636828

64. Köster J, Rahmann S. Snakemake--a scalable bioinformatics workflow engine. Bioinformatics. 2012;28: 2520–2522.

