## Supplementary Information for "A family portrait of the genomic factors shaping tandem repeat mutagenesis"

**Supplementary Information** for Sasani et al. (2026), *A family portrait of the genomic factors shaping tandem repeat mutagenesis.*

**Supplementary Figures:**

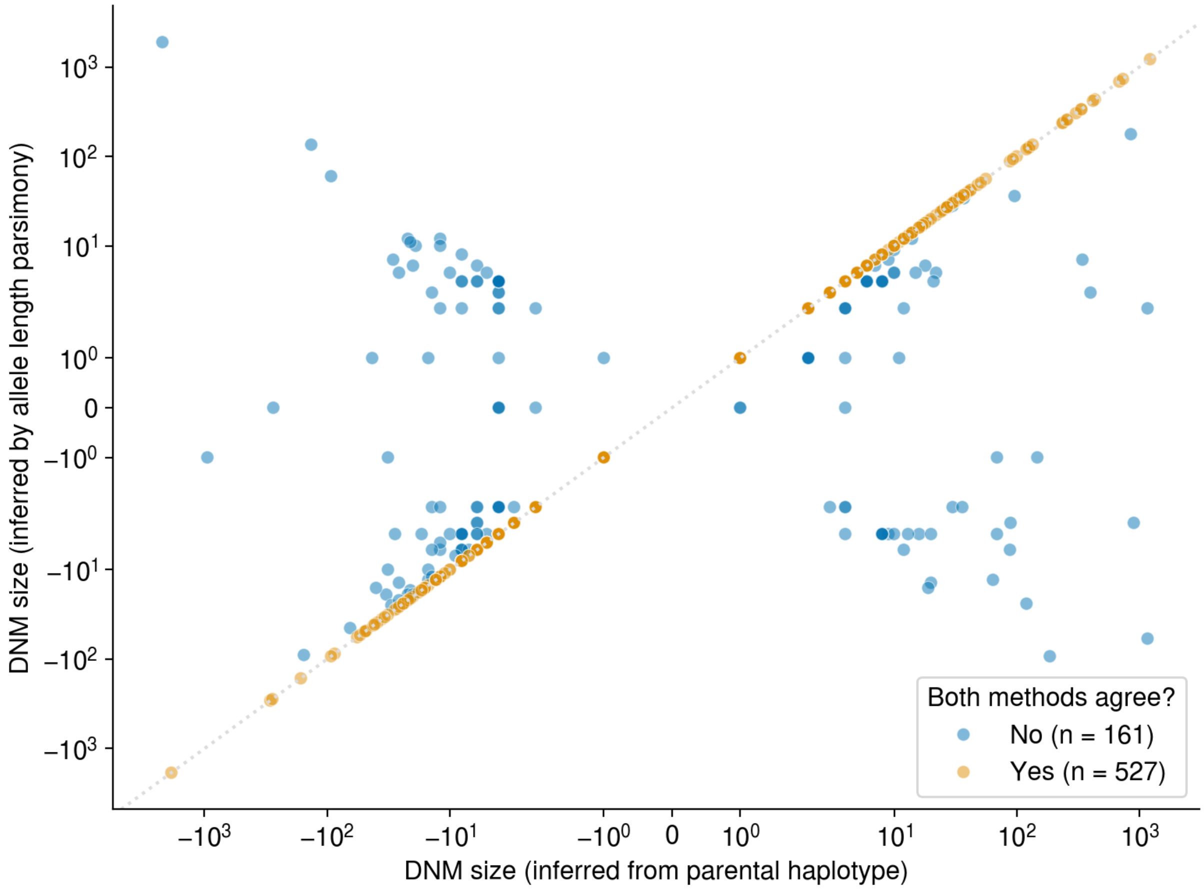


**Supplementary Figure 1**: Precisely inferring the sizes of *de novo* expansions and contractions using long reads**.** We inferred the likely size (that is, the number of base pairs that expanded or contracted to produce a *de novo* allele) at 688 *de novo* TR mutations using two methods. The first, called the "parsimony-based" method, involves comparing the *de novo* allele length in a child to the diploid allele lengths of the parent in whose germline the DNM likely occurred. The smallest of these allele length differences is assumed to represent the size of the mutational event. In the second approach, we used HiPhase [1] to identify the specific parental allele that mutated to produce the *de novo* allele, and compared the length of the *de novo* allele to the length of the "precursor" parental TR allele.

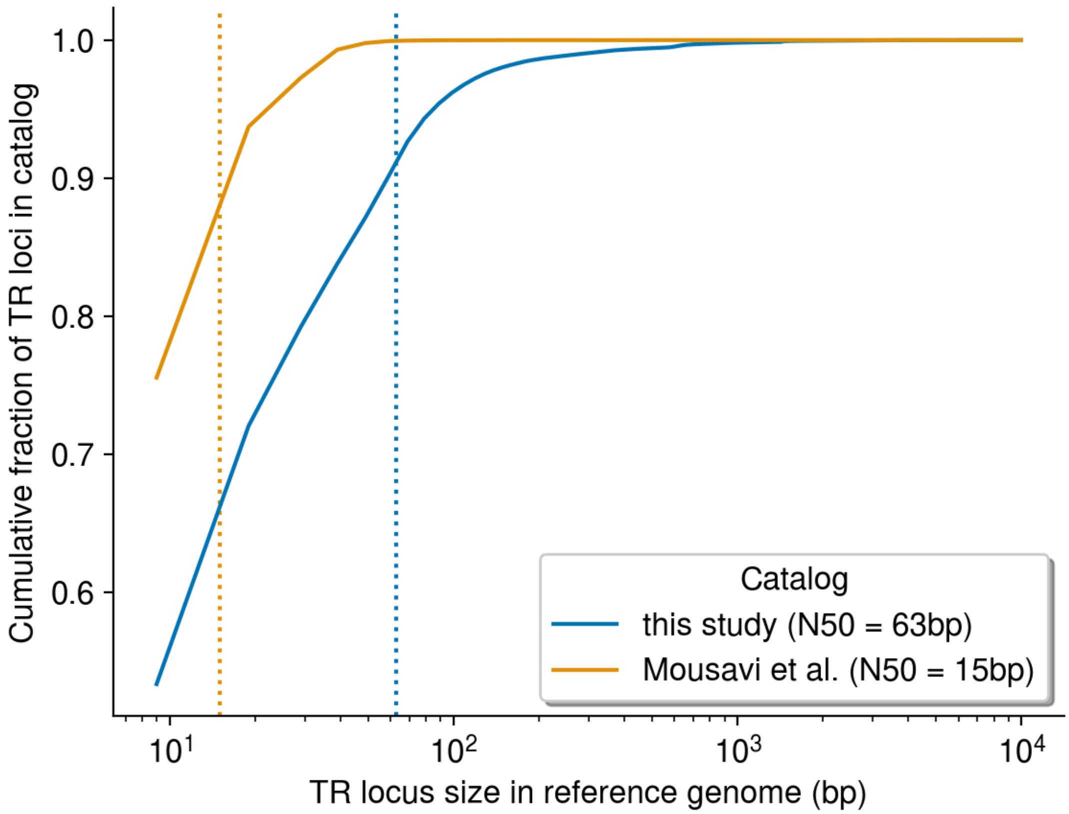


**Supplementary Figure 2**: The TR catalog used in this study is enriched for longer loci. Distributions of TR locus sizes (as measured with respect to the GRCh38 reference genome sequence) in the catalog used in [2] and the present study (shown in blue), compared to the catalog used by GangSTR [3] (shown in orange). Locus N50s (such that 50% of the TR catalog comprises loci of that size or larger) are shown as orange and blue dotted lines for Mousavi et al. (2019) and Porubsky et al. 2025, respectively.

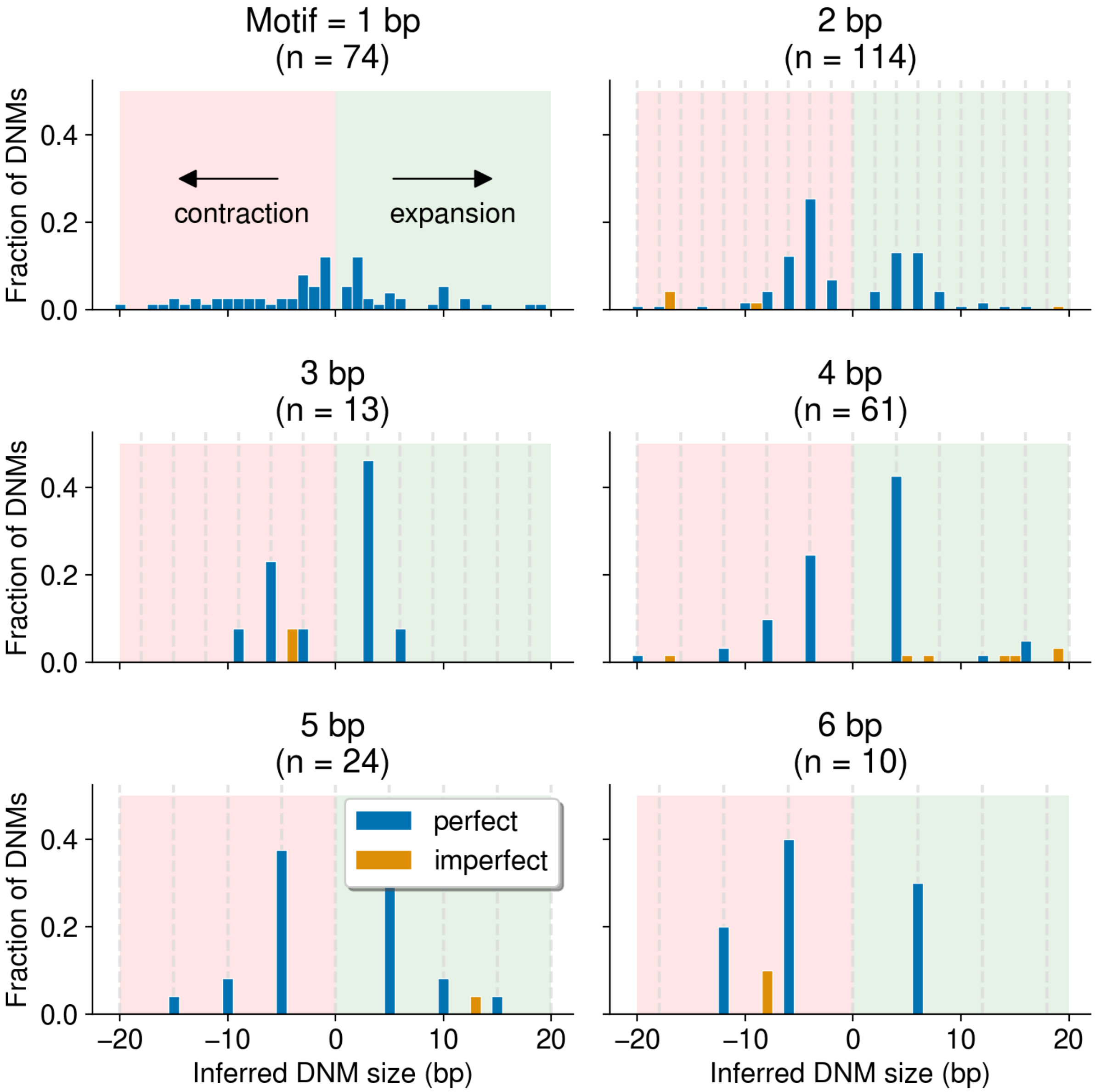


**Supplementary Figure 3**: No significant enrichment of expansions after genotyping a smaller catalog of STR loci. We genotyped a collection of approximately 1.34 million STR loci from a previous study [3] using our PacBio HiFi, TRGT, and TRGT-denovo pipeline. As in **Figure 2**, we counted the number of STR DNMs that involved an expansion or contraction of the specified number of base pairs. Blue bars correspond to counts of DNMs that were perfect multiples of the motif size, and orange bars correspond to DNMs that were imperfect multiples (e.g., a 5bp expansion of a dinucleotide motif). Dotted vertical lines indicate the expected sizes of "perfect" expansions and contractions.

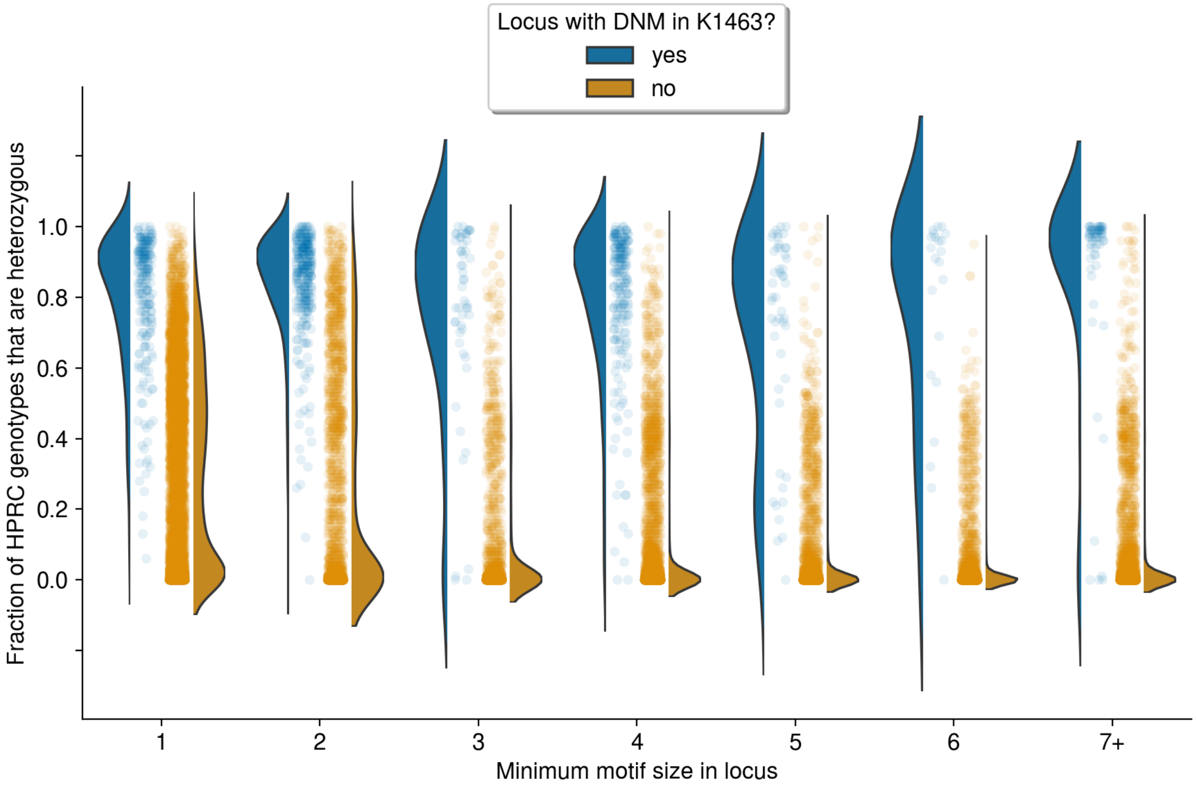


**Supplementary Figure 4:** <u>Mutable TR loci in K1463 are more heterozygous in the HPRC.</u> We calculated the number of HPRC individuals (*n* = 100) with non-identical tandem repeat allele lengths (which we refer to as "heterozygosity") at every locus at which we observed a *de novo* expansion or contraction in the K1463 pedigree (n = 1,270 loci). We then randomly sampled n = 100,000 loci from our catalog of ~7.8 million TR loci that did not mutate in K1463, and calculated HPRC heterozygosity at each locus. We then stratified loci by the size of the smallest nucleotide motif in the locus definition. Each point represents a single locus, and violin plots indicate kernel density estimates of the heterozygosity distributions.

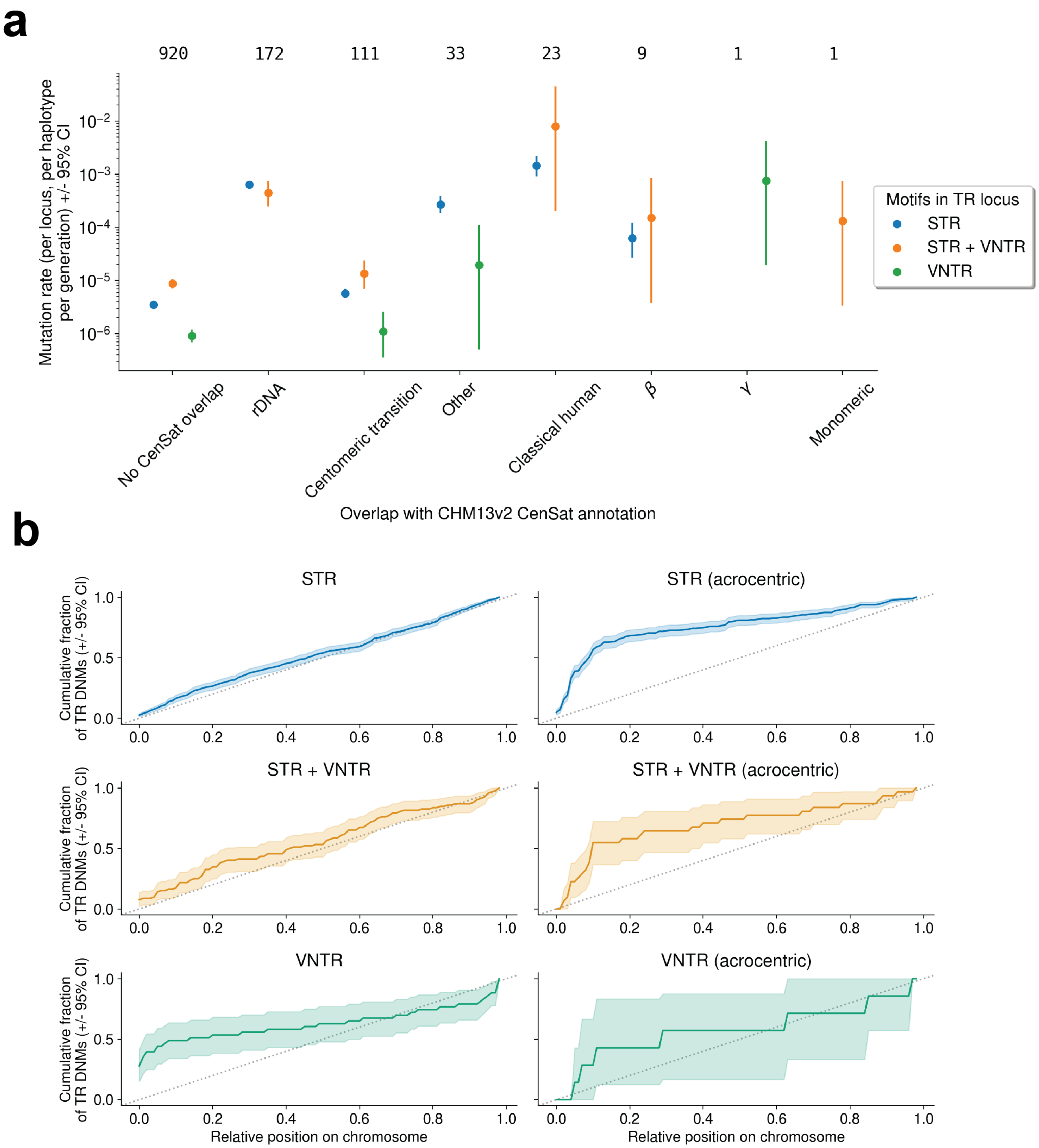


**Supplementary Figure 5:** <u>Tandem repeat *de novo* mutation rates are elevated in centromeric satellites and enriched on acrocentric chromosomes</u>. **a)** We downloaded the CenSat (hub_3671779_censat) annotations for the CHM13v2 reference genome from the UCSC Table Browser and computed the overlap between every *de novo* TR mutation in K1463 and a CenSat annotation. We then computed the overlap between every TR locus in our catalog of 7.8 million loci and the CenSat annotations. Using the loci overlapping each CenSat annotation, we then calculated the *de novo* mutation rate (expressed per locus, per haplotype, per generation) for TR loci that comprised STR motifs, VNTR motifs, or a mix of both. CenSat annotations from the CHM13v2 assembly are shown on the x-axis. The total number of TRs in each category is shown at the top of the plot. **b)** We determined the relative chromosomal position of each TR DNM (expressed as a fraction of the total chromosome length), and calculated the cumulative proportion of TR DNMs that occurred along the relative length of chromosomes in the T2T/CHM13v2 assembly (shown as a colored line in each plot). We calculated the bootstrap 95% confidence interval for each cumulative distribution, which is shown as a shaded region around the colored line.The cumulative distribution function for the uniform distribution is shown as a dotted grey line. We separately plot the cumulative proportions of TR DNMs that occurred on acrocentric chromosomes (chr13, chr14, chr15, chr21, and chr22) or non-acrocentrics.

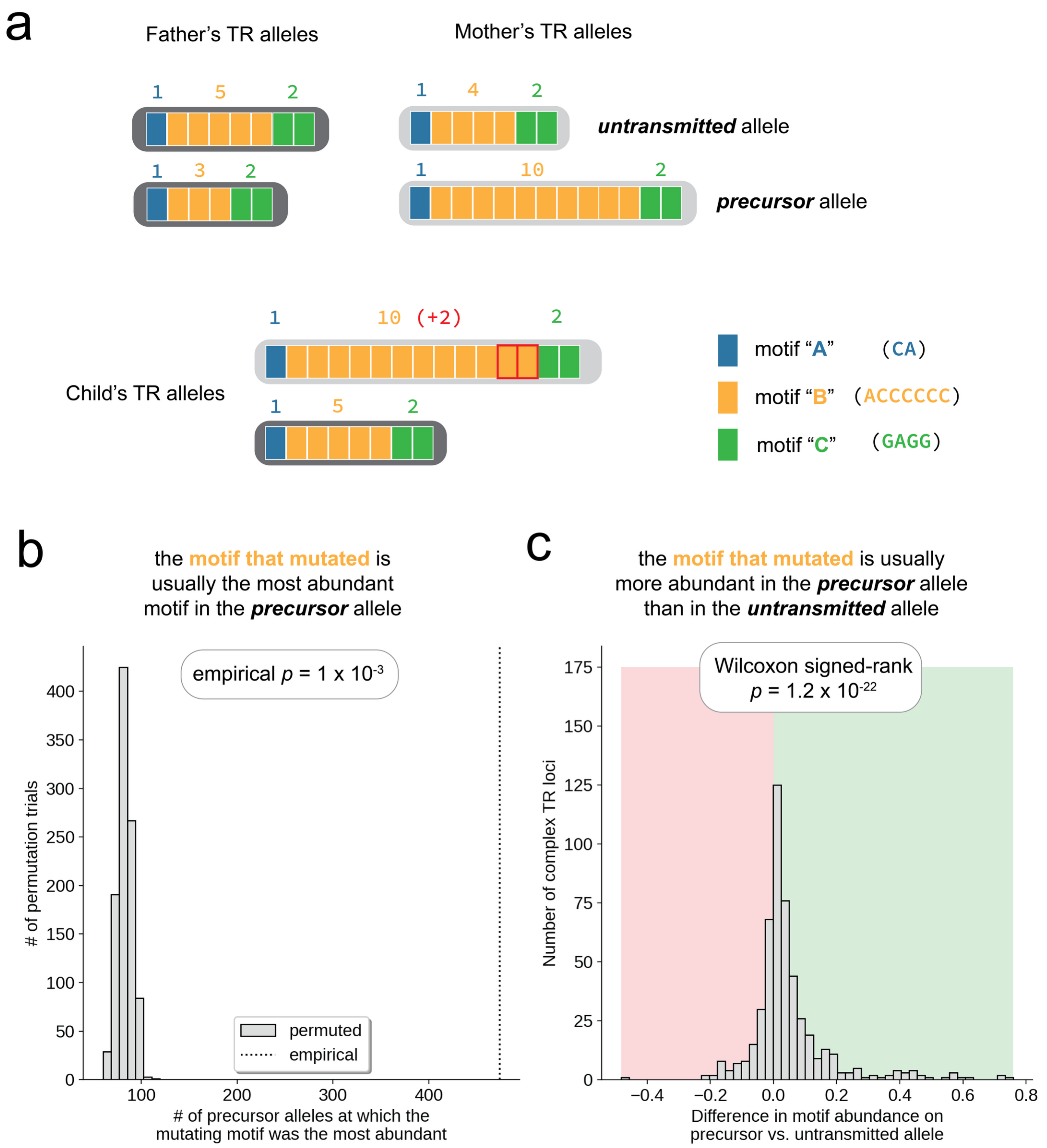


**Supplementary Figure 6:** At complex loci, motifs that mutate comprise longer arrays than those that don't mutate**. a)** Toy diagram of a tandem repeat locus, which comprises three motifs ("A," "B," and "C"). The child possesses a *de novo* allele with 12 copies of motif "B," which is the product of a 2-motif expansion of "B" motifs in the mother's germline. In the parent-of-origin, the allele that mutated is denoted the "precursor" allele, and the allele that was not inherited by the child is denoted the "untransmitted" allele. **b)** On each precursor allele, we calculated the relative abundance of each constituent motif. We then counted the number of precursor alleles at which the motif that mutated was also the most abundant motif on the allele. In each of 1,000 trials, we permuted the motif labels at each precursor allele and recalculated the number of times the “mutating” motif was the most abundant. **c)** At each complex TR locus, we compared the abundance of the motif that mutated on the precursor allele to the abundance of that motif on the untransmitted allele (positive values indicate that motif abundance was higher on the precursor allele). We then performed a paired Wilcoxon rank-sum test to compare the abundance of the mutating motif on precursor vs. untransmitted alleles.

**Supplementary Note** (including Supp. Fig. 7 – 9)**:**

*De novo TR mutation discovery is highly sensitive to sequencing depth*

We originally sequenced members of the K1463 pedigree with PacBio HiFi to an average genome-wide depth of approximately 40X, with the exception of NA12877 and NA12878 (the members of the second generation, G2), who were each sequenced to nearly 100X [2]. Because NA12877 and NA12878 were sequenced to very high depths, we focused our original analysis of *de novo* TR mutation on DNMs observed in their children (the members of the third generation, G3). When we examined *de novo* TR mutations observed in the fourth generation (G4), we noticed several unusual patterns. We identified far more *de novo* mutations in the members of G4, of which a much higher fraction occurred at homopolymers (**Supplementary Figure 7a**). We also observed a striking parent-of-origin bias among the G4 DNMs. In sub-family G4A, almost 90% of DNMs were assigned to the paternal haplotype, and in sub-family G4B, nearly 75% were assigned to the maternal haplotype (**Supp. Fig. 7b**). In both cases, we assigned more DNMs to the parent with lower sequencing depth than the other. In the absence of other compelling evidence, we hypothesized that *de novo* TR discovery and parent-of-origin inference in G4 might be biased by parental sequencing depth. To address this hypothesis, we performed a series of downsampling experiments.

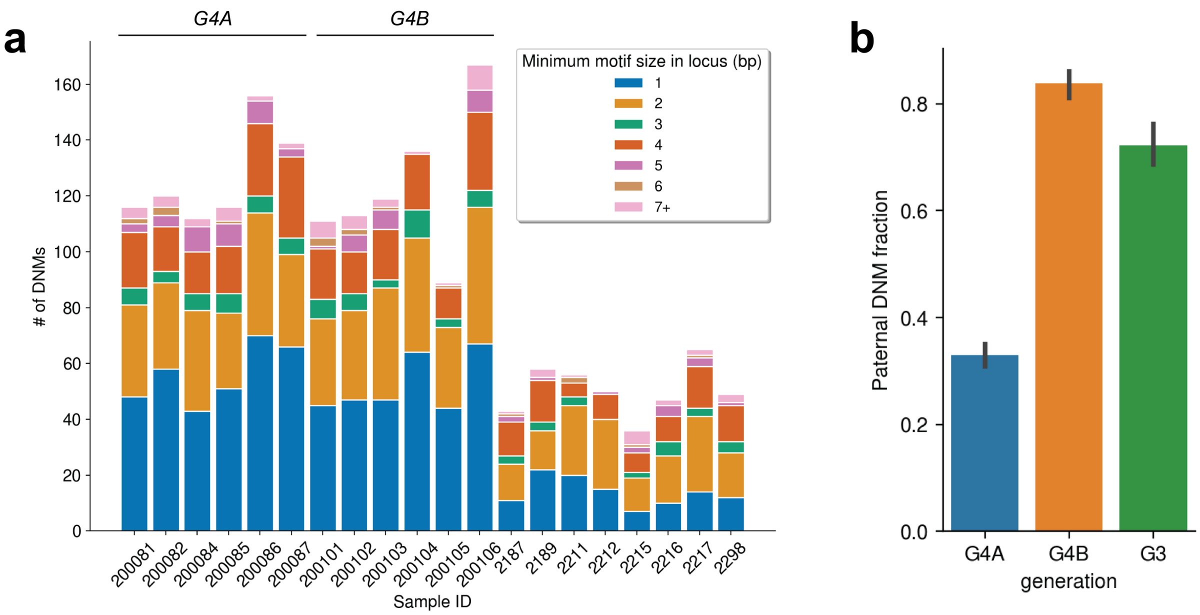


**Supplementary Figure 7**: G4 DNMs were enriched at homopolymer loci in G4 and exhibited unusual parent-of-origin biases**.** All analyses were performed using the "original" sequencing data from each member of K1463. **a)** Counts of *de novo* tandem repeat mutations in each child, stratified by the size of the smallest motif at each TR locus. **b)** Average fraction (+/- 95% CI) of *de novo* mutations assigned to the paternal gamete-of-origin in the DNMs identified in each generation.

We selected a single trio from the K1463 pedigree and performed a series of downsampling experiments with that trio's PacBio read alignments. This trio included two parents (NA12877 and NA12878) who were sequenced to nearly 100X depth and one child (NA12885) who was sequenced to approximately 80X. To investigate the impact of parental sequencing depth on TR genotyping and *de novo* expansion or contraction discovery, we first randomly downsampled every individual in the trio to 50X depth using samtools [1]. Then, we randomly downsampled either of the two parent's BAM files to 10, 20, 30 or 40X while leaving the other parent and the child's BAM files unchanged. Using these new BAM files, we genotyped all 7.8 million TR loci in each individual using TRGT [5] and searched for *de novo* TR alleles in the child using TRGT-denovo [6]. We also performed single-nucleotide variant calling in each downsampled BAM file using DeepVariant [7], and jointly phased the SNV and TR genotypes in each individual using HiPhase [1]. Finally, we used these phased genotypes to infer the likely parent-of-origin for each *de novo* TR mutation as described in the **Materials and Methods**.

Downsampled parental sequencing depth had a dramatic impact on TR genotyping and *de novo* TR discovery. When one parent had lower sequencing depth than the other, we identified far more DNMs than when both parents were sequenced to the same depth, and these excess DNMs were enriched for *de novo* mutations at homopolymers (**Supp. Fig. 8)**. We also observed a striking parent-of-origin bias in the phased DNMs (**Supp. Fig. 9**), such that we assigned many more *de novo* mutations than expected to the parent with lower sequencing depth. This is likely a result of "allelic dropout." If a TR allele is not captured in the parent's sequencing reads, but is inherited by their child, that allele will erroneously appear to be *de novo*. Imbalanced parental depths were responsible for substantial parent-of-origin biases, even when both parents were sequenced to at least 30X genome-wide coverage, a typical threshold for whole-genome sequencing studies (**Supp. Fig. 9**).

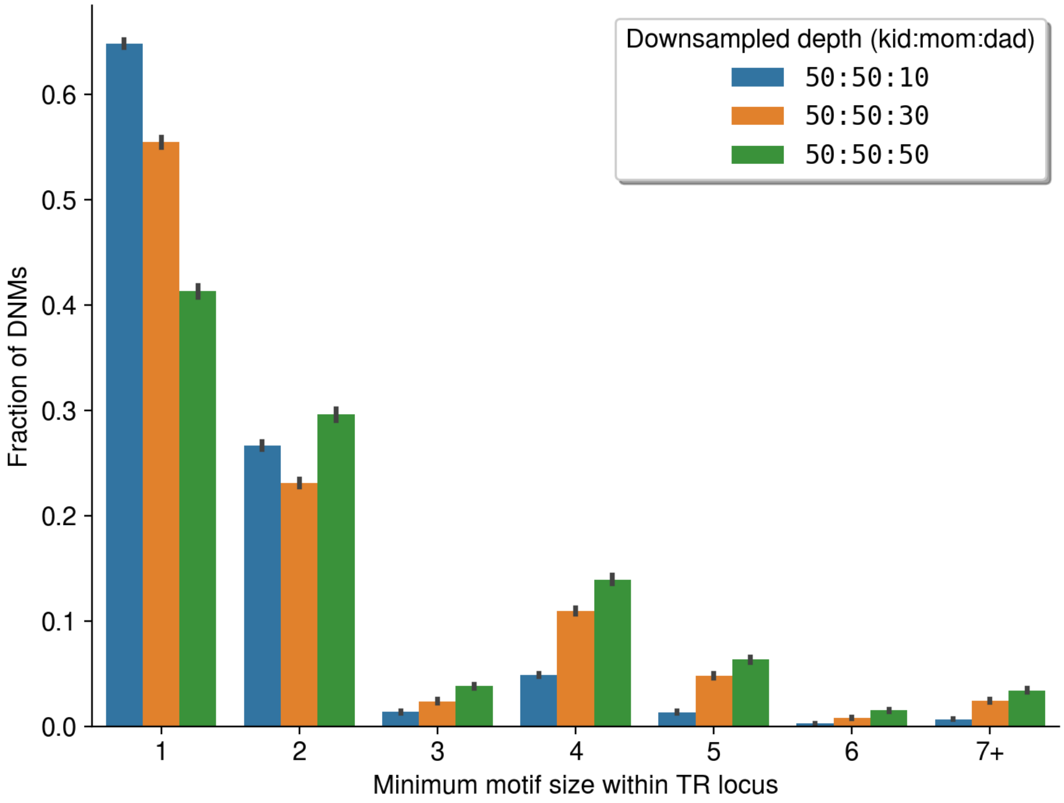


**Supplementary Figure 8**: Imbalanced parental sequencing depth has a major impact on the motif patterns of TR DNMs**.** Using a single trio (NA12877, NA12878, NA12885), we randomly downsampled the father's BAM file while leaving the other parent's and the child's BAM files at 50X. We then performed TR genotyping and *de novo* discovery, as well as small variant calling and haplotype phasing, using the updated trio BAMs. We calculated the size of the smallest motif within each TR locus, and counted the fraction of DNMs that occurred at loci with that minimum motif size.

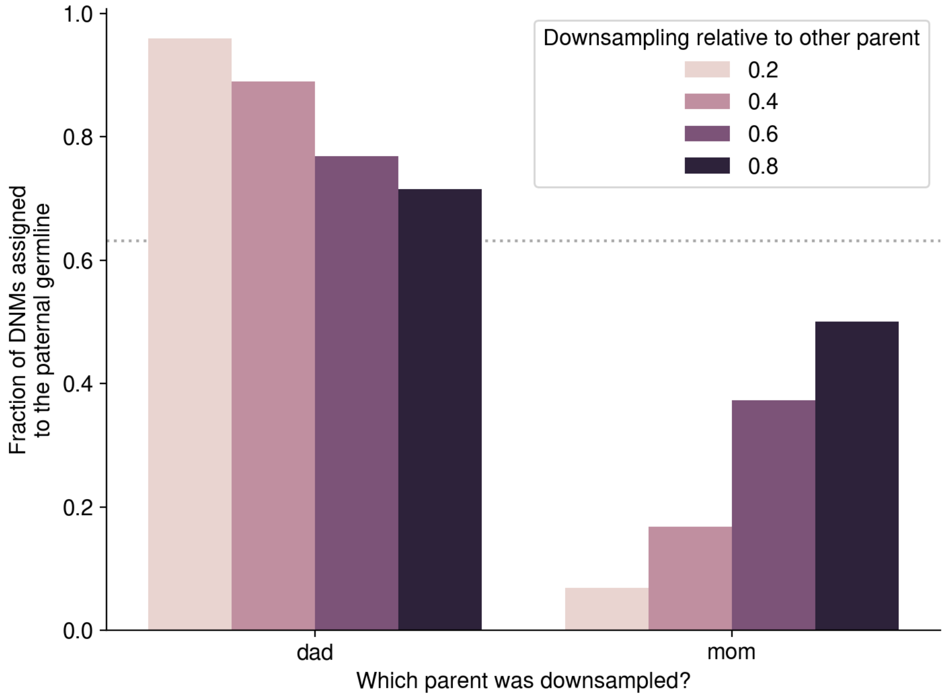


**Supplementary Figure 9**: Imbalanced parental sequencing depth has a major impact on parent-of-origin inference at TR DNMs**.** Using a single trio (NA12877, NA12878, NA12885), we randomly downsampled one parent's BAM file from 50X to either 10X, 20X, 30X, or 40X while leaving the other parent's and the child's BAM files at 50X. We then performed TR genotyping and *de novo* discovery, as well as small variant calling and haplotype phasing, using the updated trio BAMs. We then inferred the parent-of-origin for each *de novo* TR mutation. We consistently assigned a larger-than-expected fraction of DNMs to the germline of the "downsampled" parent; based on prior evidence, we expect approximately 70% of DNMs to originate in the paternal germline (shown as a dotted grey line). As parental sequencing depth became more balanced, the fraction of DNMs assigned to the paternal germline steadily approached this expectation.

Having confirmed the major impact of parental sequencing depth on *de novo* TR discovery and genotyping, we re-sequenced the four parents of the G4 family members in K1463 to a total genome-wide average of approximately 75X coverage. We then re-genotyped all members of the K1463 pedigree with the latest version of TRGT [5] and identified *de novo* TR expansions and contractions using TRGT-denovo [6] (**Materials and Methods**). As expected, using "topped-up" sequencing data in the four parents of the G4 children, we observed a comparable number of TR DNMs in G3 and G4 and the expected paternal origin ratio.

1. Holt JM, Saunders CT, Rowell WJ, Kronenberg Z, Wenger AM, Eberle M. HiPhase: jointly phasing small, structural, and tandem repeat variants from HiFi sequencing. Bioinformatics. 2024;40. doi:10.1093/bioinformatics/btae042

2. Porubsky D, Dashnow H, Sasani TA, Logsdon GA, Hallast P, Noyes MD, et al. Human de novo mutation rates from a four-generation pedigree reference. Nature. 2025;643: 427–436.

3. Mousavi N, Shleizer-Burko S, Yanicky R, Gymrek M. Profiling the genome-wide landscape of tandem repeat expansions. Nucleic Acids Res. 2019;47: e90.

4. Danecek P, Bonfield JK, Liddle J, Marshall J, Ohan V, Pollard MO, et al. Twelve years of SAMtools and BCFtools. Gigascience. 2021;10. doi:10.1093/gigascience/giab008

5. Dolzhenko E, English A, Dashnow H, De Sena Brandine G, Mokveld T, Rowell WJ, et al. Characterization and visualization of tandem repeats at genome scale. Nat Biotechnol. 2024;42: 1606–1614.

6. Mokveld T, Dolzhenko E, Dashnow H, Nicholas TJ, Sasani T, van der Sanden B, et al. TRGT-denovo: accurate detection of de novo tandem repeat mutations. bioRxivorg. 2024. doi:10.1101/2024.07.16.600745

7. Poplin R, Chang P-C, Alexander D, Schwartz S, Colthurst T, Ku A, et al. A universal SNP and small-indel variant caller using deep neural networks. Nat Biotechnol. 2018;36: 983–987.
